# mTORC2 couples fasting to mitochondrial fission

**DOI:** 10.1101/2022.07.19.500669

**Authors:** Nuria Martinez-Lopez, Pamela Mattar, Miriam Toledo, Henrietta Bains, Manu Kalyani, Marie Louise Aoun, Mridul Sharma, Laura Beth J. McIntyre, Leslie Gunther-Cummins, Frank P. Macaluso, Jennifer T. Aguilan, Simone Sidoli, Mathieu Bourdenx, Rajat Singh

## Abstract

Fasting triggers diverse cellular and metabolic adaptations to facilitate organismal survival^1,2^. During nutrient deprivation, increases in circulating fatty acids support mitochondrial respiration^2^. The mechanisms driving mitochondrial adaptations and respiratory sufficiency during nutrient deprivation remain incompletely understood. Here we show that extended periods of fasting, or lipid availability stimulates mTORC2 activity. Activation of mTORC2 and phosphorylation of its target NDRG1^3^ at S336 sustains mitochondrial fission and respiratory sufficiency. Timelapse imaging reveals that wildtype NDRG1, but not phosphorylation-deficient NDRG1^S336A^ mutant, engages with mitochondria to facilitate its scission. Using proteomics, and an siRNA screen, we show that mTORC2-phosphorylated NDRG1 cooperates with the small GTPase Cdc42^4^ and Cdc42-specific effectors and regulators to orchestrate fission. Accordingly, *Rictor*^KO^, NDRG1^S336A^ mutants, and *Cdc42*-deficient cells each display mitochondrial phenotypes reminiscent of fission failure. During nutrient surplus, mTOR complexes perform anabolic functions^5^; however, paradoxical reactivation of mTORC2 during fasting plays an unexpected role in driving mitochondrial fission and respiration.

---

Fasting increases circulating fatty acids, which are used for organismal sustenance^2^. To understand mechanisms driving metabolic adaptations during fasting or fatty acid availability, we sought to identify signaling cascades activated under these conditions. To this purpose, unbiased quantitative phosphoproteomics was performed using nanoscale liquid chromatography-mass spectrometry (nLC-MS/MS) in livers from mice that were (1) basal fed; (2) overnight (14-16 h) fasted; or gavaged with (3) dietary triglycerides as corn oil or (4) BODIPY C_16_/palmitic acid or (5) refed a high-fat diet after 14-16 h fast **(Fig. 1A)**. We incorporated corn oil or BODIPY C_16_ cohorts to compare signaling mechanisms activated by dietary exogenous fatty acids versus lipolysis-derived endogenous fatty acids during fasting. The refed group served to simulate physiological feeding. Delivery of BODIPY C_16_ to livers was confirmed by fluorescence **(Extended Data Fig. 1A)**. Phosphoproteomics in five groups identified 2,160 phosphosites across 942 phosphoproteins, of which 863 phosphosites (39.95%) were significantly modulated. Unsupervised hierarchical clustering grouped the basal and refed cohorts (groups 1 and 5) into one cluster, while lipid-exposed groups, i.e., fasted, corn oil, and BODIPY C_16_ (groups 2, 3 and 4) clustered into the second group **(Extended Data Fig. 1B)**. Clustering of phosphosites to determine phosphoproteins that are coordinately modulated revealed a major “green cluster” encompassing 86.9% of significantly modulated phosphosites **(Fig. 1B)**. Z score-normalized abundance of phosphosites in green cluster was significantly higher in fasted, corn oil, and BODIPY C_16_ groups compared to basal and refed groups **(Fig. 1C)**. Interestingly, despite strong reduction, green cluster-normalized abundance was higher in refed mice compared to basal, indicating qualitative differences in phosphopeptides between both groups **(Fig. 1C)**.

**Fig. 1.**
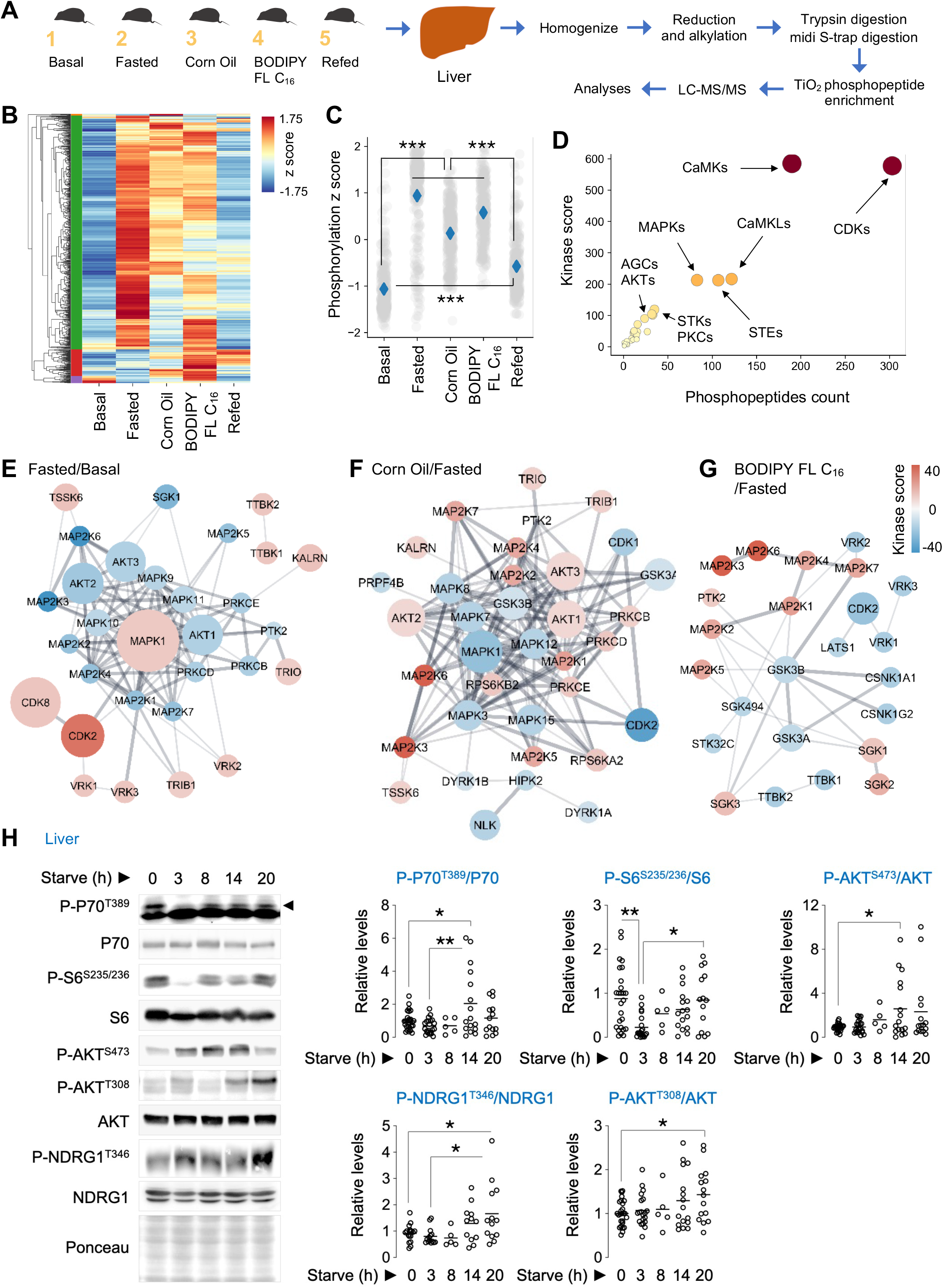
Hepatic phosphoproteome and kinases predicted to respond to fasting or lipids. **(A)** Phosphoproteomics in livers as per plan shown in cartoon (n=4). **(B)** Heatmap and hierarchical clustering of phosphosites across indicated groups indicated in **A. (C)** Phosphoproteome-wide comparisons via z score normalization of phosphosites in green cluster. Gray dots represent individual phosphosites. Blue diamonds represent group means. Non-parametric ANOVA [Kruskal-Wallis statistic=797.3, P<0.0001] followed by Dunn’s multiple comparison test, ***P<0.001 (n=4). **(D)** Prediction for kinases that putatively target the phosphosites identified within the green cluster in **B**, and **(E-G)** pairwise comparisons showing upregulated or downregulated kinase networks. **(H)** Immunoblots (IB) and quantification for indicated proteins in livers of 2-10 months (mo)-old male and female mice that were fed or fasted for the indicated durations (n=5-27 mice). Ponceau is loading control. Individual replicates and means are shown. *P<0.05 and **P<0.01, One-way ANOVA followed by Tukey’s multiple comparisons test. Please refer to Table S2_statistical summary. Please refer to excel files, proteomics1a_igps_green_cluster, and proteomics1b. h = hours

To determine kinases putatively modulating phosphosites in the green cluster, we used iGPS^6^, which revealed that these phosphosites are targets of cyclin-dependent kinases (CDKs), Ca^2+^/calmodulin-dependent kinases (CaMKs), mitogen-activated protein kinases (MAPKs), and Ser/Thr AGC kinases **(Fig. 1D, Extended Data Fig. 1C)**. Pairwise comparisons showed upregulation of CDK2/CDK8 and MAPK1 during fasting when compared to basal group **(Fig. 1E)**. Corn oil, which is devoid of cholesterol or proteins, and BODIPY C_16_, each perturbed a number of kinase groups albeit to a lesser extent when compared to fasting. Interestingly, modest enrichment of RPS6KA2/B2, AKT1-3, PRKCB/D, and SGK1-3 with corn oil, and SGK1-3 in BODIPY C_16_ group, pointed to mTORC1 or mTORC2 activation **(Fig. 1F, G)**. Refeeding reduced the overall kinase network, but expectedly activated nutrient-sensitive kinases, e.g., mTOR, and suppressed those induced by fasting, e.g., CDK2 **(Fig. 1E, Extended Data Fig. 1D)**. Given our interest in the mTOR-autophagy axis, we confirmed that corn oil exposure robustly increased levels of P-P70^T389^, P-S6^S235/236^, and P-AKT^S473^, and Raptor and Rictor—broadly reflecting mTORC1/C2 activation; without affecting the energy sensor^7^, P-AMPK^T172^ **(Extended Data Fig. 1E, F)**. Refeeding or exposure to corn oil each increased P-P70^T389^ and P-AKT^S473^ to similar levels suggesting that availability of lipids originating from breakdown of dietary triglycerides leads to mTORC1/C2 activation **(Extended Data Fig. 1E, F)**. We next asked if increased fatty acid availability during fasting **(Extended Data Fig. 1G)** associates with mTORC1/C2 reactivation as observed with corn oil. Strikingly, fasting for 14 h or longer robustly reactivated mTORC1 and mTORC2 signaling in liver **(Fig 1H)**, but not in adipose tissue **(Extended Data Fig. 2A)**, indicated by phosphorylation of their respective targets^3^, P70^T389^, and AKT^S473^ and NDRG1^T346^ **(Fig 1H)**. By contrast, fasting had no effect on phosphorylated levels of PKA^T197^, PKCα/βII^T638/641^, PKCδ^T505^, PKCδ/θ^S643/676^ and PKCζ/λ^T410/403^ in liver **(Extended Data Fig. 2B)**—indicating specificity for fasting-induced reactivation of the AKT and SGK1 arms, but not PKA/C cascades, of mTORC2 signaling. Whether reactivation of mTOR during fasting in vivo is due to lipid availability or increased lysosomal proteolysis as previously described^8^, remains untested. Although insulin regulates mTOR activity, mTOR activation in response to corn oil appears to not require insulin since corn oil does not elicit a substantial insulin release **(Extended Data Fig. 3A-E)**, and because equivalent mTORC1/C2 activation was observed in control and streptozotocin (STZ)-injected mice, which reduces insulin levels and causes hyperglycemia due to β-cell destruction **(Extended Data Fig. 3A-E)**. Thus, fasting or lipid availability each stimulates mTORC1/C2.

To determine physiological roles of mTOR reactivation during fasting, we inactivated mTORC1 or mTORC2 or hyperactivated mTORC1 by knocking out respective regulatory genes *Raptor, Rictor* or *Tsc1* using liver-restricted AAV8-TBG-Cre **(Extended Data Fig. 4A)**. Loss of mTORC2 activity in liver, and not adipose tissue or muscle, was confirmed by markedly reduced AKT^S473^ phosphorylation **(Extended Data Fig. 4B, C)**. We examined the effect of loss of each gene on liver triglyceride (TG) levels and respiration rates. While control and mTORC1 inactivated (*Raptor*^KO^) livers showed equivalent liver TGs during fasting, hyperactivation of mTORC1 (*Tsc1*^KO^) lowered liver TGs **(Extended Data Fig. 4D)** consistent with mTORC1’s role in VLDL secretion^9^. Surprisingly, in contrast to the established TG lowering effect of *Rictor* loss in fed/obesogenic states^10^, inactivating mTORC2 (*Rictor*^KO^) markedly increased liver TGs and lipid droplet content during fasting **(Fig. 2A, Extended Data Fig. 4E)** without affecting circulating free fatty acids **(Extended Data Fig. 4F)**. Interestingly, *Rictor*^KO^ livers showed lower oxygen consumption rates **(Fig. 2B)** and accumulation of substrates for mitochondrial respiration, acyl carnitines **(Fig. 2C)**, which in conjunction with reduced mitochondrial membrane potential in *Rictor* depleted cells **(Extended Data Fig. 4G)** indicated mitochondrial insufficiency. Decreased oxygen consumption was not due to impaired expression of fatty acid oxidation or electron transport genes—in fact, fasted *Rictor*^KO^ livers displayed increased expression of fat oxidation (*Ppara, Cpt1a, Cpt1b, Cact, Cpt2*), mitochondrial biogenesis (*Ppargc1a*), and electron transport (*Cox1-4, Nd1-3, Cytb, Atp8, Atp6*) genes **(Extended Data Fig. 5A-P)**. In addition, protein levels of mitochondrial fatty acid uptake proteins (CPT1a/CPT2/CACT) and electron transport chain components, NDUFB8 (complex I), SDHB (II), UQCRC2 (III), MT-CO1 (IV), and ATP5A (V) were comparable in control and *Rictor*^KO^ livers **(Extended Data Fig. 5V, W)**. Interestingly, loss of *Rictor* led to increased expression of mitochondrial fusion genes *Mfn1/2* and *Opa1* during fasting without affecting fission genes *Mff* and *Dnm1l* (Drp1) **(Extended Data Fig. 5Q-U)**—directing us to formally investigate whether reactivation of the broadly anabolic mTORC2 during fasting paradoxically supports mitochondrial dynamics and respiration.

**Fig. 2.**
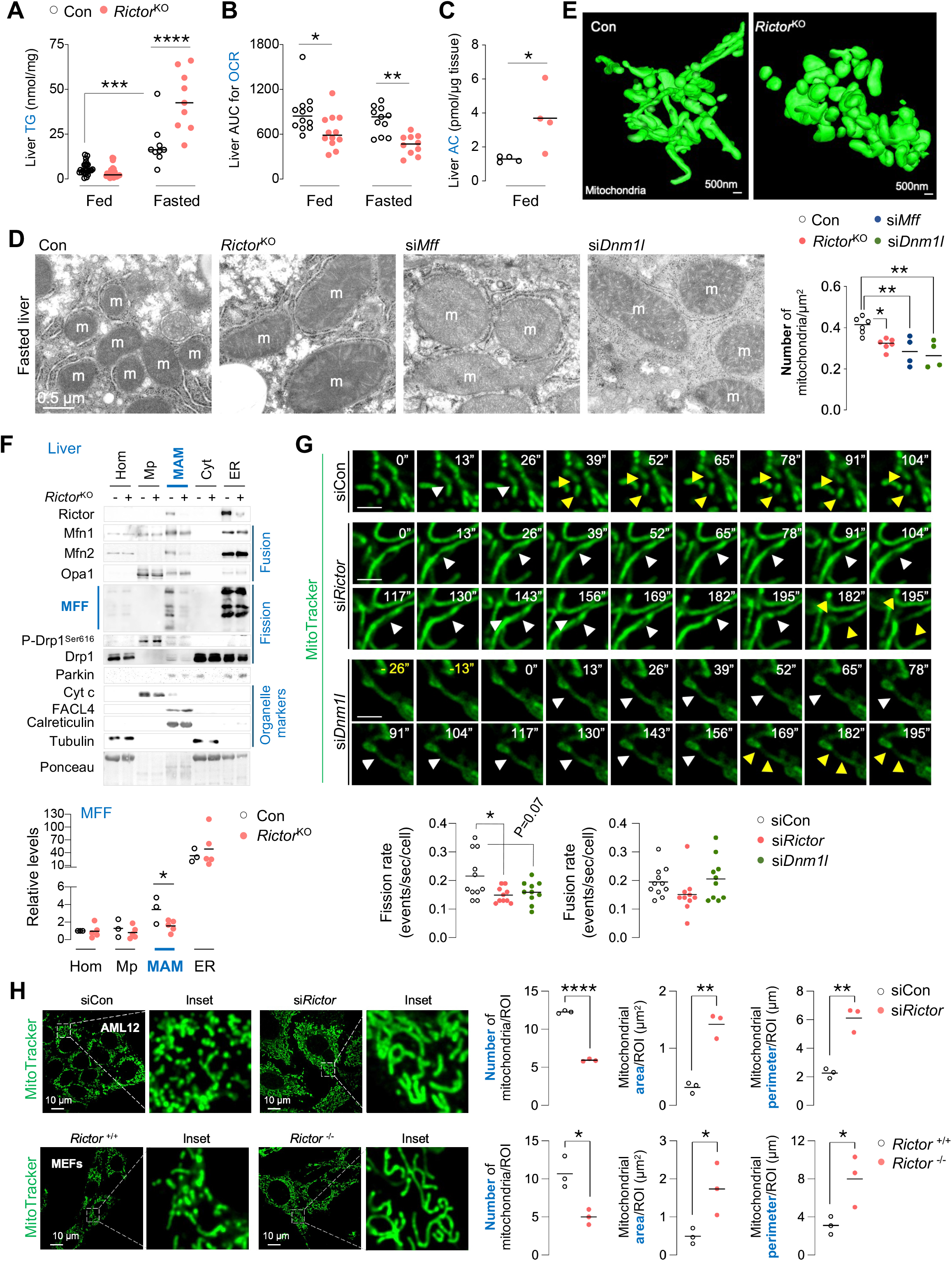
Loss of mTORC2 dampens mitochondrial fission and respiratory function. **(A)** Liver triglycerides (TG) (n=9-24), and **(B)** area under curve (AUC) for oxygen consumption rates (OCR) in Con and *Rictor*^KO^ livers from fed or 14-16 h fasted mice (n=10-12). **(C)** Acylcarnitine (AC) content in livers of 4-5 mo-old Con or *Rictor*^KO^ male mice (n=4). **(D)** Conventional electron micrographs (EM) in 14-16 h fasted Con, *Rictor*^KO^, si*Mff* or si*Dnm1l* livers of 4-9 mo-old male mice (n=4-6 mice). Quantification for mitochondrial number is shown. **(E)** Three-dimensional reconstruction of electron micrographs of mitochondria (green) from Con or *Rictor*^KO^ livers. **(F)** IB and quantification for indicated proteins in homogenates (Hom), pure mitochondria (Mp), mitochondria-associated membranes (MAM), cytosol (Cyt), and endoplasmic reticulum (ER) fractions from 14-16 h fasted Con and *Rictor*^KO^ livers (n=3-5). Two livers were pooled to generate 1 sample. **(G)** Live cell imaging and quantification for mitochondrial fission and fusion rates in siCon, si*Rictor* or si*Dnm1l* NIH3T3 cells (n=10-11 cells from n=6 independent experiments). Scale = 2 µm. **(H)** Representative confocal images of **(top)** AML12 cells cells knocked-down for *Rictor*, and **(bottom)** *Rictor*^*-/-*^ MEFs and their corresponding controls cultured in serum-free medium for 30 min in presence of MitoTracker green to stain mitochondria. Magnified insets are shown. Quantification for mitochondrial number, area and perimeter is shown (AML12 = 45 cells; MEFs = 36-43 cells from n=3 independent experiments each). Individual replicates and means are shown. *P<0.05, **P<0.01, ***P<0.001 and ****P<0.0001, 2-way ANOVA followed by Tukey’s multiple comparison test **(A** and **B)**; One-way ANOVA followed by Tukey’s multiple comparisons test **(D** and **G)**; unpaired Student’s t-test **(C, F** and **H)**. Please refer to Table S2_statistical summary.

To determine if mTORC2 plays a role in mitochondrial dynamics during fasting, we first asked how fasting impacts mitochondrial dynamics *in vivo*. Electron microscopy (EM) analyses of fasted livers revealed increased mitochondrial number with greater frequency of smaller mitochondria (with reduced area, perimeter, and length)—independent of changes in mitochondrial mass (equivalent VDAL1 and Cyt c levels), when compared to fed livers—reflecting increased mitochondrial fission during fasting **(Extended Data Fig. 6A, B)**. By contrast, *Rictor*^KO^ livers and those silenced for fission genes, *Mff*^11^ or *Dnm1l*^12^ **(Extended Data Fig. 6C)**, each failed to show an increase in mitochondrial number during fasting **(Fig. 2D)**, in turn, leading to increases in mitochondrial area and perimeter **(Fig. 2D, Extended Data Fig. 6D, E)**—reflecting fission failure during fasting. Correlating with impaired fission in *Rictor*^KO^ livers, 3D EMs revealed broad mitochondrial defects including mitochondrial distention and blunted inter-mitochondrial networking **(Fig. 2E)**, while TEM showed reduced mitochondrial-ER contacts^13^ during fasting **(Extended Data Fig. 6F)**. *Rictor*^KO^ livers exhibited mild changes in lipid composition of mitochondrial associated membranes (MAMs)^13,14^ (as determined via unbiased lipidomics) and no global increases in markers of ER stress or proteostasis failure, excluding their direct contribution to the mitochondrial phenotype **(Extended Data Fig. 7B, C)**. Because Rictor/mTORC2 localizes to ER-mitochondrial contact sites/MAMs **(Fig. 2F)** to regulate Ca^2+^ homeostasis and apoptosis^13^, we envisioned that mTORC2’s role at MAMs extends to the regulation of mitochondrial fission. Consistent with this idea, MAMs were enriched in Rictor, and in proteins regulating fission^15^ (mitochondrial fission factor (MFF) and Drp1) and fusion (Opa1 and mitofusins) **(Fig. 2F)**. Strikingly, MAMs from *Rictor*^KO^ livers showed marked reduction in MFF levels without affecting levels of P-Drp1^Ser616^, Drp1, mitofusins or Opa1 **(Fig. 2F)**—supporting the idea that loss of mTORC2 blocks mitochondrial fission.

To confirm mTORC2’s role in mitochondrial fission, we used time-lapse microscopy to assess changes in rates of mitochondrial fission and fusion in si*Rictor* cells in real time. This endeavor revealed that loss of fission in si*Rictor* cells was comparable to fission failure in cells knocked down for a *bona fide* driver of fission, *Dnm1l* **(Fig. 2G)**. Since fusion rates were identical in siCon and si*Rictor* cells, the observed mitochondrial phenotype in *Rictor* depleted cells is due to impaired fission and not excessive fusion. Finally, we used two hepatocyte lines, AML12 cells and HepG2 cells, silenced for *Rictor*, as well as *Rictor*^-/-^ mouse fibroblasts to determine the impact of inactivated mTORC2 signaling on fission **(Fig. 2H, Extended Data Fig. 6G-I)**. As expected, all three mTORC2 inactivated cells showed decreased mitochondrial number and marked increases in mitochondrial area and perimeter—demonstrating mTORC2’s role in stimulating mitochondrial fission. Interestingly, loss of fission in *Rictor*^-/-^ mouse fibroblasts occurred independent of changes in levels and S616 phosphorylation of Drp1 (**Extended Data Fig. 6G)**, a modification that stimulates fission^16^. On this basis, we propose that mTORC2 reactivation during fasting stimulates mitochondrial fission.

Since mTORC2 phosphorylates its targets via AGC kinases^17^, AKT, PKC, and SGK1, we hypothesized that phosphorylated downstream targets of mTORC2 support mitochondrial fission during fasting. Quantitative nLC-MS/MS in control and *Rictor*^KO^ livers **(Fig. 3A)** identified 4,553 phosphosites from 1,712 phosphoproteins, of which 309 phosphosites (145 upregulated and 164 downregulated) (6.79%) on 212 phosphoproteins (12.38%) were significantly modulated in *Rictor*^KO^ livers **(Extended Data Fig. 8A)**. Gene ontology (GO) analysis of differentially modulated phosphosites followed by enrichment map network analysis^18^, allowing clustering of molecular functions based on similarity within GO terms, identified clusters related to *cytoskeleton and cellular architecture*, which are known to be regulated by mTORC2, as well as *mRNA processing and splicing, protein targeting* and *regulation of cellular catabolic processes* **(Fig. 3B, Table S1)**, and all these clusters were associated with proteins with decreased phosphorylation status. The denser cluster populated by both upregulated and downregulated phosphoproteins contained terms *regulation of metabolism*. Since protein function is modulated by site-specific phosphorylation or cumulative phosphorylation of multiple phosphosites^19^, we measured the overall phosphorylation status (ΔPs) of phosphoproteins quantified in our dataset, which revealed that phosphorylation was significantly increased (hyperphosphorylation, ΔPs > 2α) in 41 phosphoproteins and decreased in 50 phosphoproteins (hypophosphorylation, ΔPs < -2σ) **(Fig. 3C)**. Interestingly, when looking through hypophosphorylated protein targets, phosphorylation of the mitophagy adapter BNIP3 at Ser(S)79 and S88 was significantly reduced in *Rictor*^KO^ livers **(Fig. 3D, Extended Data Fig. 8B)**, with no known physiological roles assigned to BNIP3 S79/S88 phosphorylation. We also focused on NDRG1 **(Fig. 3E)**, a protein involved in lipid metabolism^20^ and extensively phosphorylated on its C terminus by mTORC2/SGK1^21^. Although NDRG1 only showed trends towards hypophosphorylation in total liver homogenates **(Fig. 3E, Extended Data Fig. 8C)**, phosphoproteomics in *Rictor*^KO^ MAMs from livers of 14-16 h fasted mice showed marked NDRG1^S336^ hypophosphorylation when compared to fasted control MAMs **(Extended Data Fig. 8D-F)**. Since mTORC2-driven NDRG1^T346^ phosphorylation increases in fasted livers (**Fig. 1h**) and because NDRG1 is present in MAMs **(Extended Data Fig. 8G)**, which are cellular hubs for regulation of mitochondrial dynamics, we sought to confirm if NDRG1^S336^ is indeed an mTORC2 target by performing phosphoproteomics of FLAG-NDRG1 pulled-down from siCon and si*Rictor* cells. To that end, the extracted ion chromatogram of peptide SRTASGSSVTS(ph)LEGTRSR, corresponding to FLAG-NDRG1 from siCon cells or si*Rictor* cells, and its relative quantification **(Extended Data Fig. 8H-J)** showed that this phosphopeptide is markedly less enriched in si*Rictor* cells compared to siCon cells—confirming that mTORC2 phosphorylates NDRG1^S336^ *in vitro* and *in vivo* in MAMs.

**Fig. 3.**
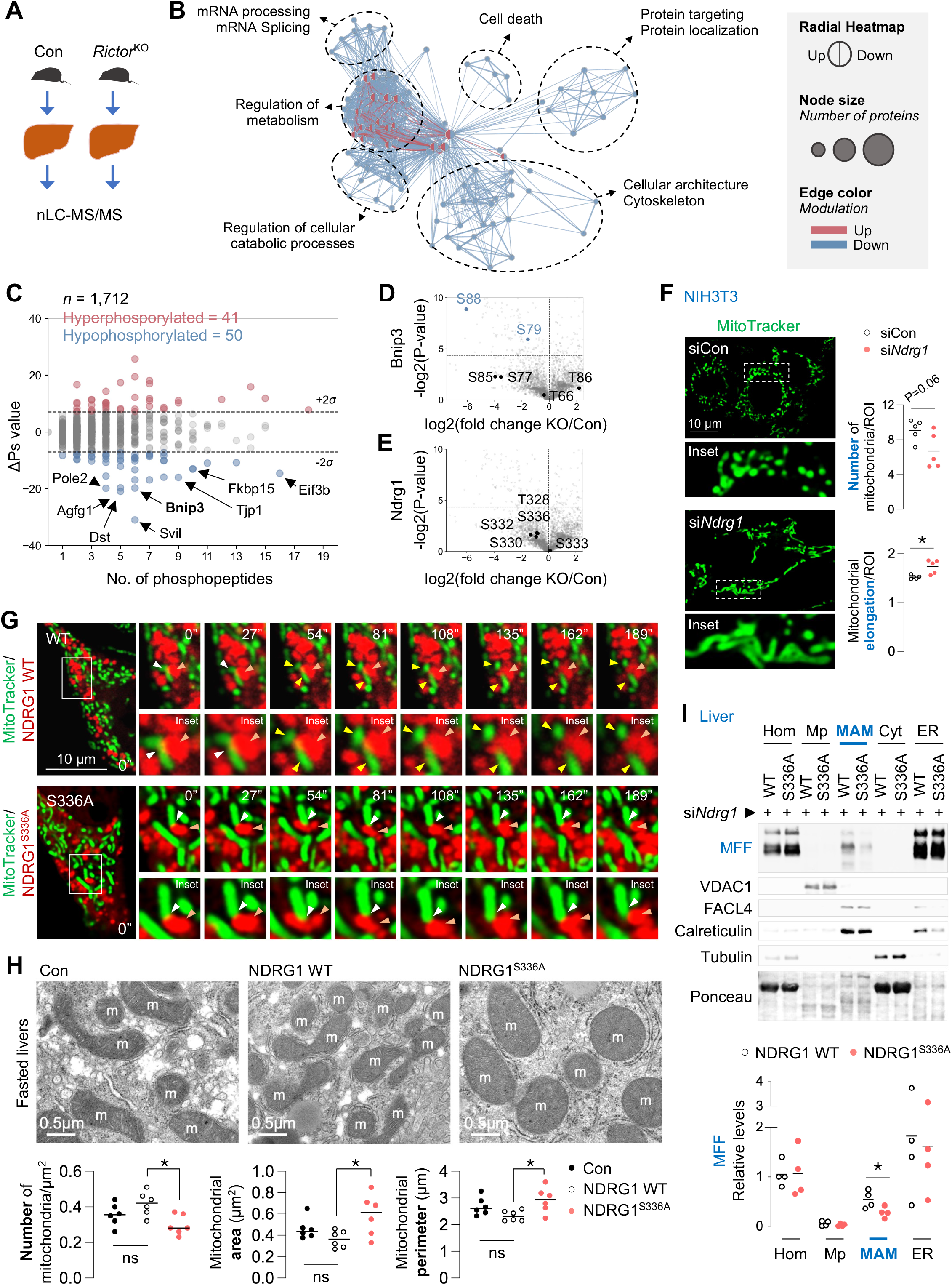
mTORC2-driven NDRG1^Ser336^ phosphorylation mediates mitochondrial fission. **(A)** Experimental plan for **B-E. (B)** Enrichment map-based network visualization of gene ontology enrichment for differentially modulated phosphosites. Blue edges show similarity between decreased phosphosites, and red nodes show similarity between increased phosphosites. Node size indicates the number of proteins per node; major clusters are circled. Associated name represents the major functional association. **(C)** Global ΔPs analyses of phosphoproteins. Hyperphosphorylated and hypophosphorylated peptides in each comparison are shown. Labels indicate the genes encoding the proteins. Dotted lines: ΔPs = ±2σ. **(D, E)** Volcano plot for **(D)** Bnip3 or **(E)** NDRG1 phosphorylation in *Rictor*^KO^ vs. Con livers fasted for 14-16 h. For **A-E** (n=3). **(F)** Representative MitoTracker green fluorescence in serum-starved siCon and si*Ndrg1* NIH3T3 cells (84-91 cells from 5 independent experiments). Quantifications for mitochondrial number and elongation are shown. **(G)** Representative live-cell images of mCherry-NDRG1 WT or mCherry-NDRG1^S336A^ in serum-starved NIH3T3 cells in presence of MitoTracker green for 30 min. Orange arrowheads: NDRG1. White arrowheads: NDRG1-mitochondria contact prior to fission. Yellow arrowheads: divided mitochondria after fission by NDRG1. Magnified insets are shown. **(H)** EM with mitochondrial quantifications (n=6 mice), and **(I)** IB and quantification for indicated proteins in Hom, Mp, MAM, Cyt, and ER fractions from livers of 3-4 mo-old NDRG1 WT or NDRG1^S336A^ male mice injected with siRNA against endogenous *Ndrg1* and fasted for 14-16 h (n=3-4 mice). Ponceau is loading control. Individual replicates and means are shown. *P<0.05. Unpaired Student’s t-test **(F** and **I)**; One-way ANOVA and Tukey’s comparison test **(H)**. ns = not significant. Please refer to Table S2_statistical summary, and Table S1_Enrichment_map. Please refer to excel file, Proteomics2_Liver_PhosphoSites.

To determine if mTORC2 mediates its effects on mitochondria by phosphorylating BNIP3 at S79 or S88 or NDRG1 at T328, S332 or S336, we set up in vitro *Seahorse*-based mito stress screens. To that purpose, we expressed phosphorylation-deficient mutant forms of FLAG-tagged BNIP3 or NDRG1 and validated equivalent FLAG expression by western blotting **(Extended Data Fig. 9A, B)**. We expressed mutant BNIP3 or NDRG1 wherein selected S or T residues on each protein were converted to alanine (A) or phosphomimetic forms of BNIP3 or NDRG1 by mutating S or T residues to aspartate (D). Serum-starved and oleic acid (OA)-treated cells (to emulate fasting in vitro) expressing WT BNIP3 or mutant S79A or S88A BNIP3 showed equivalent mitochondrial respiration **(Extended Data Fig. 9C)**—eliminating a role of BNIP3^S79/S88^ phosphorylation in mitochondrial respiration. Expressing NDRG1 T328A or S332A phosphorylation-deficient mutants or T328D or S332D phosphomimetics also failed to impact mitochondrial respiration **(Extended Data Fig. 9D, E** and **G)**. By contrast, blocking NDRG1^S336^ phosphorylation significantly reduced basal and maximal respiration and ATP production **(Extended Data Fig. 9F, G)**, while phosphomimetic NDRG1^S336D^ stimulated mitochondrial respiration compared to WT NDRG1 **(Extended Data Fig. 9F, H)**. Furthermore, silencing *Ndrg1* **(Extended Data Fig. 9I, J)** or expressing each phosphorylation-deficient NDRG1 mutant **(Extended Data Fig. 9K)** substantially lowered mitochondrial membrane potential—suggesting that NDRG1^S336^ phosphorylation mediates mTORC2’s role in regulating mitochondrial function—thus setting the basis to investigate if mTORC2 drives mitochondrial fission by NDRG1^S336^ phosphorylation. Several S/T residues on the NDRG1 C-terminus are phosphorylated by SGK1. In support for the idea that mTORC2/SGK1 axis mediates fission via NDRG1^S336^ phosphorylation, we noted that silencing *Sgk1-3* or *Ndrg1*, but not *Akt1/2*, significantly reduced mitochondria number and increased mitochondrial area, perimeter, and elongation **(Extended Data 10A)**, as typically observed in cells silenced for *Dnm1l*^22^ or *Mff*^23^. Consistent with these findings in cultured cells, silencing *Sgk1* and *Ndrg1*, and not *Akt1/2*, reduced cellular respiration *in vivo* **(Extended Data 10B, C)**.

To determine if mTORC2-driven NDRG1^S336^ phosphorylation facilitates mitochondrial fission, we used time-lapse microscopy to first determine if and why NDRG1 interacts with mitochondria. Interestingly, NDRG1 WT frequently engages with mitochondria in what appears to be colocalization with a constricted region of mitochondria—culminating in fission **(Fig 3G, Extended Data 11A, E)**. Quantification for duration of NDRG1-mitochondrial interactions, and whether each interaction leads to mitochondrial fission (useful) or not (futile), revealed that WT-NDRG1-mitochondrial interactions resulted in fission within ∼90.8+/-11.1s of contact **(Extended Data 11E, F)**. By contrast, mutant NDRG1^S336A^ maintained its colocalization with ER **(Extended Data 12A, B)** but exhibited futile interactions with mitochondria that were in average 289.8+/- 26.7s long and did not cause fission **(Fig 3G, Extended Data 11B, E, F)**. As expected, knocking-down *Rictor* also prolonged the duration of interaction of WT NDRG1 with mitochondria (409.5+/-63.7s) and blocked the ability of WT NDRG1 to divide mitochondria **(Extended Data 11C, E, F)**—linking mTORC2-driven NDRG1^S336^ phosphorylation to mitochondrial fission. Strikingly, knocking-down *Dnm1l* failed to block WT NDRG1’s ability to divide mitochondria **(Extended Data 11D-F)**, at least in this in vitro model, reflected by ∼102+/-24.6s taken for NDRG1 to divide mitochondria under *Dnm1l* depleted conditions. These data suggest that while Drp1 is a key regulator of fission, it appears to not influence mTORC2-NDRG1 regulation of mitochondrial fission.

To determine whether phosphorylation-deficient mutant NDRG1^S336A^ in livers recapitulates the mitochondrial phenotype of *Rictor*^KO^ livers, we expressed FLAG-tagged NDRG1 WT or NDRG1^S336A^ in livers silenced for endogenous *Ndrg1* **(Extended Data Fig. 12C)** and confirmed equivalent FLAG expression by immunohistochemistry **(Extended Data Fig. 12D)**. Consistent with our observations in *Rictor*^KO^ livers, fasted NDRG1^S336A^ livers showed enlarged mitochondria with reduced mitochondrial number, and increased area and perimeter **(Fig 3H)**—reflecting impaired fission. As observed in *Rictor*^KO^ livers, fasted NDRG1^S336A^ livers also showed reduced cellular respiration **(Extended Data Fig. 12E)**. Furthermore, MAMs from *Rictor*^KO^ and NDRG1^S336A^ livers, both showed lower levels of mitochondrial fission protein MFF when compared to corresponding controls **(Fig 2F, 3I)**. Despite reduced levels of MFF in MAMs from *Rictor*^KO^ livers, levels of total and S616 phosphorylated Drp1, which drive fission^16^ remained unaffected in *Rictor*^KO^ MAMs **(Fig. 2F)**. Taken together, these in vitro and in vivo data show that mTORC2 stimulates mitochondrial fission by phosphorylating NDRG1 at S336.

Since NDRG1 does not exhibit intrinsic GTPase activity, a critical requirement for membrane scission, we asked whether P-NDRG1^S336^ engages with proteins with intrinsic GTPase activity to facilitate fission. To identify proteins bound to NDRG1 that could serve this function, we used proteomics to find proteins bound to FLAG NDRG1 WT but not NDRG1^S336A^. Intriguingly, we found that NDRG1 WT interacts with Cdc42, a Rho GTPase that regulates actin cytoskeleton^24^ and cytokinesis^25^ **(Extended Data Fig. 13A)**. Indeed, when compared to NDRG1 WT, NDRG1^S336A^ displayed modestly reduced binding to Cdc42, Arhgef10 (a Rho GEF that activates Rho GTPases by stimulating GDP/GTP exchange), and Arhgap35 (Rho GAP that facilitates GTP hydrolysis to inactivate Rho GTPases) likely due to the transient and dynamic nature of these interactions. This led to our hypothesis that Cdc42 mediates the effects of mTORC2-NDRG1^Ser336^ in mitochondrial fission.

To begin to test this hypothesis, we performed co-IP experiments, which confirmed that exogenously expressed GFP-Cdc42 interacts with FLAG-NDRG1 WT but fails to interact with mutant NDRG1^S336A^ **(Extended Data Fig. 13B)**. Furthermore, FLAG-NDRG1 WT interacts with mCherry-Cdc42 WT, but displays reduced interaction with mutant Cdc42^T17N^, which does not bind GTP **(Extended Data Fig. 13C)**— indicating that NDRG1^S336^ phosphorylation and Cdc42-GTP binding are required for their interaction to promote fission. Consistent with this idea, knocking-down *Cdc42* blocked NDRG1’s ability to drive fission under serum starved conditions **(Extended Data Fig. 13D, E)**. In addition, while mCherry-Cdc42 WT engaged with, and divided, mitochondria in ∼157 ± 26 seconds **(Fig 4D, Extended Data Fig. 13G, H)**, mCherry-Cdc42^T17N^ exhibited prolonged (∼339 ± 58 seconds) interactions with mitochondria that did not lead to fission **(Extended Data Fig. 13F-H)**—indicating that Cdc42 GTP-binding and NDRG1^S336^ phosphorylation are both required for mitochondrial division. In support of the role of the mTORC2-NDRG1-Cdc42 axis in mitochondrial fission, knocking-down *Rictor* or *Ndrg1* or *Cdc42* each resulted in reduced mitochondrial number and increased mitochondrial area, perimeter and elongation **(Fig. 4A)**— recapitulating mitochondrial fission failure observed in cells knocked-down for *bona fide* fission factors, *Mff* or *Dnm1l*—and contrasting against fusion failure phenotypes, i.e., smaller mitochondrial size and greater mitochondrial number, noted in *Opa1* silenced cells **(Fig. 4A)**. In keeping with this, silencing *Cdc42* impaired mitochondrial function reflected by reduced mitochondria membrane potential **(Extended Data Fig. 13I)**.

**Fig. 4.**
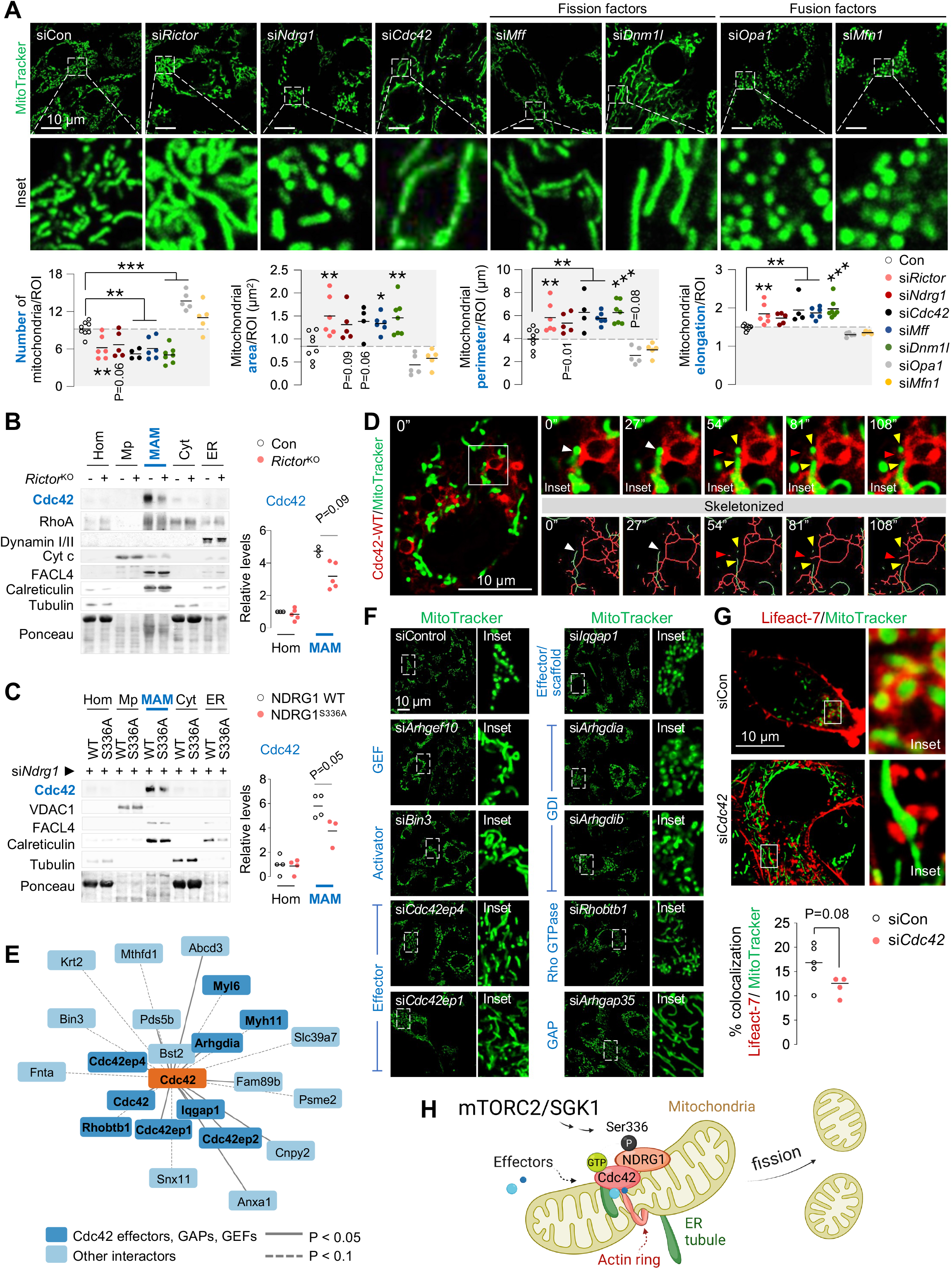
mTORC2-NDRG1-Cdc42 axis stimulates mitochondrial fission. **(A)** Representative confocal images of NIH3T3 cells transfected with indicated siRNAs and cultured in serum-free medium with MitoTracker green for 30 min. Magnified insets are shown. Quantification for mitochondria number and morphology parameters is shown. (64-142 cells from n= 5-8 independent experiments). Gray areas indicate mitochondria fission-deficient models. **(B, C)** IB and quantification for indicated proteins in Hom, Mp, MAMs, Cyt and ER from livers of 14-16 h-fasted **(B)** 5-6 mo-old control and *Ricto*r^KO^ (n=3-5 mice), or **(C)** 3-4 mo-old mice expressing NDRG1 WT or NDRG1^S336A^ plasmids co-injected with siRNA against endogenous *Ndrg1* (n=3-4 mice). **(D)** Representative live-cell imaging in cells expressing mCherry-tagged Cdc42 WT cultured in serum-free medium for 30 min in presence of MitoTracker green. Red arrowheads: Cdc42. White arrowheads: Cdc42-mitochondria contact prior to fission. Yellow arrowheads: divided mitochondria after contact with Cdc42. Magnified insets are shown. **(E)** Cartoon representing significantly (P<0.05) enriched interacting partners of Cdc42 in bold identified via proteomics, some of which belong to the Rho family of GTPases. **(F)** Representative images of NIH3T3 cells knocked-down for indicated Cdc42-binding partners and cultured in serum-free medium for 30 min in presence of MitoTracker green. Magnified insets are shown (n=37-40 cells from 3 independent experiments). **(G)** Representative confocal images of siCon or si*Cdc42* NIH3T3 cells expressing mCherry-Lifeact-7 cultured in serum-free medium for 30 min in presence of MitoTracker green. Magnified insets are shown. Quantification for % colocalization of mCherry with mitochondria is shown (n=30-33 cells from 4-5 independent experiments). **(H)** Reactivation of mTORC2 during fasting phosphorylates NDRG1 at Ser336, which engages with mitochondria and recruits Cdc42 to mitochondria-ER contact sites wherein Cdc42 and its effector proteins orchestrate mitochondrial fission. Ponceau is loading control. Individual replicates and means are shown. *P<0.05, **P<0.01 and ***P<0.001. One-way ANOVA and Dunnett’s multiple comparisons test **(A)**; unpaired Student’s t-test **(B, C** and **G)**. Please refer to Table S2_statistical summary. Please refer to excel files, Proteomics6_GFP_Cdc42_coIP.

Since Cdc42 is largely controlled through translocation, we hypothesized that mTORC2-driven NDRG1^S336^ phosphorylation is a signal to recruit Cdc42 to MAMs to drive fission. Indeed, Cdc42 and RhoA^26^ **(Fig 4B)**, but not dynamins, were enriched in MAMs from fasted control livers. By contrast, MAMs from fasted *Rictor*^KO^ **(Fig 4B)** and NDRG1^S336A^ livers **(Fig 4C)** each showed markedly reduced Cdc42 levels without affecting levels of RhoA—pointing to a role of the mTORC2-NDRG1^S336^ axis in recruiting Cdc42 to MAMs. Given the enrichment of Cdc42 in MAMs in an mTORC2- and NDRG1^S336^-sensitive manner, it is likely that Cdc42 governs local downstream signaling controlling mitochondrial fission. Consequently, we used proteomics to identify Cdc42 regulators and effectors that may potentially regulate fission. Using NIH3T3 cells expressing GFP-Cdc42 or GFP-empty vector as negative control and analyzing fold-change interaction (using cut-off P-value of 4.32), we short-listed 11 proteins that were significantly enriched in GFP-Cdc42 pulldowns when compared to empty vector **(Fig 4E, Extended Data Fig 14A, B)**. Of note, 5 enriched targets specifically belonged to the Cdc42 family of effectors and regulators: Cdc42 effector proteins (Cdc42ep4, Cdc42ep1, and Cdc42ep2), the Rho GDP Dissociation Inhibitor alpha (Arhgdia), and IQ motif-containing GTPase-activating protein 1 (Iqgap1), which is a downstream Cdc42 effector and an upstream Cdc42 scaffold protein **(Fig 4E, Extended Data Fig 14B)**. We also observed enrichment (albeit insignificant) in GFP-Cdc42 pulldowns of a known driver of Cdc42 in muscle cells, bridging integrator 3 (Bin3), the atypical Rho GTPase/Rho GDP-dissociation inhibitor 1 (Rhotbt1), and Rho GTPase inhibitor, Rho GDP Dissociation Inhibitor (GDI) beta (Arhgdib) **(Fig 4E, Extended Data Fig 14B)**. In addition, we included two identified binding partners of NDRG1, Arhgef10 and Arhgap35, which serve as GEF and GAP, respectively **(Extended Data Fig 13A)**.

To test the role of the identified candidates in mitochondrial fission, we transfected NIH3T3 cells with siRNAs against each selected target, except *Cdc42ep2* since silencing it severely reduced viability **(Extended Data Fig 14C)**. Our results from the screen provide strong support for the role of Cdc42 and its family of effectors/regulators in mitochondrial fission, since deleting *Cdc42* or Cdc42 activators, *Arhgef10* and *Bin3* or Cdc42 downstream effectors, *Cdc42ep4* or *Cdc42ep1*, each resulted in increased mitochondrial area, perimeter and elongation—reflecting impaired fission **(Fig 4F, Extended Data Fig 14D-G)**. In addition, silencing the Rho GDP dissociation inhibitor *Arhgdia* (but not *Arhgdib*), which releases Cdc42 from inhibition by Arhgdia, increased mitochondrial number—reflecting increased fission; further supporting the role of Cdc42 in mitochondrial fission **(Fig 4F, Extended Data Fig 14D-G)**. Interestingly, not all Rho GTPases impact mitochondrial dynamics, since depletion of Rho GTPase family *Rhobtb1* gene had no effect on mitochondrial morphology, while knocking down *Iqgap1*, an effector and activator of Cdc42, increased mitochondrial number likely because Iqgap1 serves as both downstream Cdc42 effector and upstream Cdc42 scaffold protein **(Fig 4F, Extended Data Fig 14D-G)**. Since Arhgap35 inactivates GTPases, we anticipated that depleting *Arhgap35* would stimulate Cdc42, leading to fission; however, knocking down A*rhgap35* decreased mitochondrial number—reflecting fission failure **(Fig 4F, Extended Data Fig 14D-G)**. This likely reflects the complex regulation of Cdc42 requiring subsequent inactivation to complete its function^27,28^, as well as specificity amongst the different effectors and regulators in stimulating fission during fasting. Consistent with these findings, in addition to Cdc42, we also observed substantial enrichment in MAMs of Cdc42 effector, Cdc42ep1, and those of regulator proteins, Arhgap35 and Arhgdia **(Extended Data Fig 15A, B)**. Interestingly, levels of Arhgap35, Cdc42ep1 and Arhgdia in MAMs from *Rictor*^KO^ **(Extended Data Fig 15A)** and NDRG1^S336A^ expressing livers **(Extended Data Fig 15B)** were comparable to those in corresponding controls, indicating that fission is regulated at the level of recruitment of Cdc42 to MAMs. Since the actin cytoskeleton drives mitochondrial division^29^, we asked whether Cdc42 mediates the effect of mTORC2-NDRG1^Ser336^ on mitochondrial fission by remodeling the actin cytoskeleton. Indeed, in control cells, actin assembled around mitochondrial subpopulations to generate ring-like structures and promote fission^30^ **(Fig 4G)**. By contrast, silencing *Cdc42* led to decreased colocalization of actin with mitochondria, which correlated with impaired mitochondrial fission **(Fig 4G)**— supporting our notion that Cdc42 facilitates the organization of actin around mitochondria to enable fission.

In sum, we show that mTORC2, an anabolic signaling module that typically responds to nutrients and growth factors, is reactivated by fasting. Our data suggest that mTORC2-mediated NDRG1^S336^ phosphorylation allows it to serve as an adapter to recruit regulatory proteins to MAMs, which are key sites for ER-mediated mitochondrial fission. We demonstrate that NDRG1 associates with mitochondrial constrictions to facilitate fission **(Fig 4H)**, and that fission events are blocked in cells expressing the phosphorylation-deficient NDRG1^S336A^ mutant or cells lacking *Rictor* or *Cdc42* or the identified Cdc42 effector/regulators **(Fig 4H)**—revealing the existence of an mTORC2-NDRG1-Cdc42 axis to support mitochondrial fission during fasting. Given the importance of mitochondrial dynamics and respiratory sufficiency in a host of clinical scenarios and ageing, identifying targets to modulate mitochondrial dynamics in these conditions provides translatable relevance to our findings.

## Data availability

Raw files of nLC-MS/MS acquisition are uploaded in the public database Chorus (https://chorusproject.org) under Project ID #1713. The Python code required to process proteomic data has been deposited on GitHub (link to be provided upon acceptance).

## Acknowledgements

This work was supported by RF1AG043517, R01DK123327, R01AG065985 and P01AG031782 to R.S. S.S. and J.A. are supported by LRF Hollis Brownstein New Investigator Research Grant, AFAR Sagol Network GerOmic Award and NIH P30 CA013330 47. We thank the Einstein Analytical Imaging Facility, which is supported by NCI cancer center support grant P30CA013330 and SIG #1S10OD016214-01A1. We thank the Biomarkers Core Laboratory at the Irving Institute for Clinical and Translational Research, Columbia University, for lipidomic assessments. We thank Dr. Katarzyna Kulej (Childrens’ Hospital of Philadelphia) for assistance with phosphoproteomic analyses. We thank Dr. Susmita Kaushik (Albert Einstein College of Medicine) for helpful suggestions for live cell imaging. I thank Eva for support during the COVID-19 pandemic.

## Author contributions

Conceptualization, R.S., N.M.-L.; Methodology, R.S., N.M.-L.; Investigation, N.M.-L., P.M., M.T.; Writing – Original Draft, R.S.; Revised Draft, R.S., N.M.-L.; Data Analyses, R.S., N.M.-L., S.S., M.B.; Funding Acquisition, R.S.; Resources, L.C., F.M., L.B.M, J.A., S.S.; Supervision, R.S.

## Methods

### Animal models

C57BL/6 (000664), Rictor^flox/flox^ (020649), Raptor^flox/flox^ (013188) and Tsc1^flox/flox^ (005680) mice were purchased from Jackson Laboratory. Studies were performed in 2-10-month-old male and female mice. Mice were fed a regular chow diet (5058; Lab Diet, St Louis, MO, USA) and maintained at 22-23 °C under a 12h:12h light/dark cycle in the institutional barrier facility. Liver-specific *Rictor*^KO^, *Raptor*^KO^ or *Tsc1*^KO^ mice were generated by injecting the flox/flox mice retro-orbitally with 2×10^11^ genome copies (gc) of AAV8-TBG-iCre adenovirus (Vector Biolabs, VB1724) per mouse and mice were humanely-killed after 8 weeks^31^. AAV8-TBG-eGFP-injected mice were controls (Vector Biolabs, VB1743). Mice were subjected to fasting/starvation (Stv) for the period indicated in figure legends with free access to drinking water and were compared to ad-libitum fed mice. As indicated in Fig. 1, Stv mice were subjected to three treatments: oral gavage of (i) corn oil (400 μl; Sigma-Aldrich, 8267), (ii) BODIPY FL C_16_ (10 mg/kg; Invitrogen, D3821) or (iii) refed high fat diet (HFD) (60% kcal in fat; Research Diets, Inc., D12492) for 30 min. Streptozotocin (STZ) to deplete insulin required administration of low doses of STZ (40 mg/kg; Sigma-Aldrich, S0130) injected intraperitoneally once-a-day for 5 consecutive days. The protocol was repeated 8 weeks after the first injection and tissues were collected 2 weeks after the last injection. For studies comprising insulin stimulation, 14 h-fasted mice were subjected to an intraperitoneal injection of insulin (4.5 mg/kg; Sigma-Aldrich, I1882) and killed 30 min later as described^13^. Mice were used under a protocol approved by the Institutional Animal Care and Use Committee.

### Corn oil composition

(Sigma-Aldrich, 8267): Corn oil is protein-free and cholesterol-free. Corn oil upon oral gavage and subsequent digestion, provides fatty acids and monoacylglycerols, which are absorbed by the gut and delivered systemically. The composition of corn oil as purchased from Sigma-Aldrich is: Water Content ≤ 0.1%, Brassica sterol ≤ 0.3%, Fatty Acid (< C14) ≤ 0.1%, Fatty Acid (C14) ≤ 0.1%, Fatty Acid (C16) 8.6 - 16.5%, Fatty Acid (C16:1) ≤ 0.5 %, Fatty Acid (C18) 1.0 - 3.3%, Fatty Acid (C18:1) 20.0 - 42.2%, Fatty Acid (C18:2) 39.4 - 62.0 %, Fatty Acid (C18:3) 0.5 - 1.5%, Fatty Acid (C20) ≤ 0.8%, Fatty Acid (C20:1) ≤ 0.5%, Fatty Acid (C22) ≤ 0.3%, Fatty Acid (C22:1) ≤ 0.1 %, Fatty Acid (C24) ≤ 0.4%.

### Cell culture

NIH3T3 cells (ATCC, CRL-1658) and HepG2 cells (ATCC, HB-8065) were cultured in high-glucose (4.5 g/L) DMEM (Gibco, 11965118) supplemented with 10% (v/v) fetal bovine serum (FBS) (Sigma-Aldrich, 12106C) and 1% (v/v) penicillin-streptomycin (P/S) (Gibco, 15140). AML12 cells (ATCC, CRL-2254) were cultured in DMEM nutrient mixture F-12 (Gibco, 11320033) supplemented with 10% FBS, 1% Insulin-Transferrin-Selenium (Gibco, 41400-045), 40ng/ml dexamethasone (Sigma-Aldrich, D4902) and 1% P/S. Cells we maintained at 37 °C in 5% CO_2_. Wherever indicated, NIH3T3 cells were washed twice with PBS and incubated in serum-free DMEM/P/S in presence of 0.25 mM oleic acid (Sigma-Aldrich, O3008) for indicated durations. For the induction of hypoxic stress, cells exposed to the hypoxia mimetic CoCl_2_ (RICCA, 2210-4) for 24 hours as described^32^.

### Primary mouse embryonic fibroblasts (MEFs) culture

Isolation of embryonic fibroblasts from *Rictor*^flox/flox^ mice was performed as described^33^. For adenoviral infection, *Rictor*^flox/flox^ cells were plated at ∼80% confluency and infected with 50 multiplicity of infection (MOI) of adenoviral-null (Ad(RGD)-fLuc (Vector Biolabs, 9999) or Ad(RGD)-CMV-iCre (Vector Biolabs, 1769) in serum-free media for 24 h. The virus-containing media was replaced by 10% FBS medium and 72 hours post-infection cells were used for experiments.

### Generation of plasmid DNAs

Mouse NDRG1_OMu19504D, BNIP3_OMu13517D or Cdc42_OMu16203C_cDNA expression plasmids were synthesized by a commercial entity (GenScript USA Inc). NDRG1_OMu19504D and BNIP3_OMu13517D were each cloned into a pcDNA3.1+/C-(K)-DYK vector. Cdc42_OMu16203C was cloned into a pcDNA3.1(+)-N-eGFP vector. cDNA encoding NDRG1^T328A^, NDRG1^T328D^, NDRG1^S332A^, NDRG1^S332D^, NDRG1^S336A^, NDRG1^S336D^, BNIP3^S79A^ or BNIP3^S88A^_pcDNA3.1+/C-(K)-DYK mutants were generated by site-directed mutagenesis. pcDNA3.1+/C- (K)-DYK or pcDNA3.1(+)-N-eGFP vectors were use as negative controls. For live-cell imaging, mouse NDRG1_OMu19504D wild type (WT), mutant NDRG1^S336A^ or Cdc42_OMu16203C WT plasmid DNAs were cloned into a pcDNA3.1(+)-mCherry vector. cDNA encoding Cdc42^T17N^ pcDNA3.1(+)-mCherry mutant was generated by site-directed mutagenesis. mCherry-Lifeact-7 was a gift from Michael Davidson (Addgene, 54491).

### In vitro transfections of plasmid DNAs or small interfering RNA (siRNA)

In vitro transfections were performed using Lipofectamine3000 (Invitrogen, L3000) as per manufacturer’s instructions. For expression of DNA plasmids, 1.2 × 10^5^ NIH3T3 cells per ml of growth medium were transfected with 1 μg of DNA and plated in a 12-well plate dish for 48 h. For acute gene silencing, 1.2 × 10^5^ cells per ml of growth medium were transfected with the corresponding siRNA. The siRNA sequences and dose are detailed in **Table S3**. Scrambled RNA was used as negative control (siCon). Silencing efficiency was confirmed by Western blotting or qPCR for the corresponding protein or gene product, respectively.

### In vivo delivery of plasmid DNAs or siRNA

In vivo DNA delivery was performed using in vivo-jetPEI® (Polyplus-transfection® SA, 201-50G) as per manufacturer’s instructions. Briefly, 100 μg of WT NDRG1 or mutant NDRG1S336A_pcDNA3.1+/C-(K)-DYK were diluted in glucose solution and combined with 7 μl of in vivo-jetPEI® for complexation for 15 min at room temperature. 200 μl of transfection mix were administered retro-orbitally to C57BL/6 mice in a single injection 24 h prior to tissue collection. Transfection efficiency was determined by immunohistochemistry in liver tissues. Livers from non-transfected mice were used as negative controls. In vivo siRNA delivery was performed using Invivofectamine™ 3.0 Reagent (Invitrogen™, IVF3005) as per manufacturer’s instructions. Briefly, 50 μg of siRNAs against *Ndrg1, Mff* or *Drp1* were mixed with complexation buffer, added to Invivofectamine™ 3.0 Reagent in a 1:1 ration and incubated for 30 min at 50 °C. The mix was diluted in PBS pH7.4 and 200 μl of siRNA mix were administered retro-orbitally to C57BL/6 mice every 24 h for 3 consecutive days prior to tissue collection.

### RNA isolation and real-time PCR

mRNA expression was determined as described previously^34^ using M-MLV Reverse Transcriptase (Invitrogen, 28025). The primers are detailed in **Table S4**.

### Western Blotting

Total cell lysates from cells in culture were prepared using lysis buffer containing 20mM Tris pH 7.5, 50 mM NaCl, 0.5%, 1 mM EDTA, 1 mM EGTA and 1% Triton X-100 supplemented with complete EDTA-free protease inhibitor (Roche, 11873580001) and phosphatase inhibitor cocktails 2 and 3 (Sigma-Aldrich, P5726, P0044). Total protein from liver or epididymal white adipose tissue (eWAT) was isolated in RIPA buffer containing 50 mM Tris pH 8.0, 150 mM NaCl, 0.5% sodium deoxycholate, 1% SDS and 1% NP-40 supplemented with protease/phosphatase inhibitors as described above. Total protein from soleus muscle was isolated as described^35^. Lysates were centrifuged at 17,000 *g* for 30 min at 4 °C and supernatants were subjected to immunoblotting by denaturing 20-30 μg of protein at 95 °C for 5 min in 3X Laemmli sample buffer containing 187.5 mM Tris, 6% SDS, 30% glycerol, 0.03% bromophenol blue and 10% β-mercaptoethanol. For analysis of relative protein levels of OXPHOS complexes, samples were boiled at 50 °C for 5 min according to manufacturer’s protocol (Abcam, ab110413). Samples were resolved by SDS-PAGE using standard methods described previously^36^. Protein bands from immunoblots were quantified by ImageJ software (NIH) and normalized to Ponceau S. Antibodies used are detailed in **Table S5**.

### Subcellular fractionation

Freshly harvested livers were fractionated for isolation of mitochondria-associated membranes (MAM), pure mitochondria (Mp), cytosol (Cyt) and endoplasmic reticulum (ER) fractions as described^14^. Cytochrome c (Cyt c) and Voltage Dependent Anion Channel 1 (VDAC1) were enrichment markers for mitochondria; long chain fatty acid-coenzyme A ligase 4 (FACL4), an enrichment marker for MAM; calreticulin as marker for MAMs and ER, while tubulin was as purity control to exclude cytoplasmic contamination.

### Co-Immunoprecipitation (Co-IP)

For the pulldown assays performed in **Extended data Fig 13B**, total cell lysates (1000 μg) from NIH3T3 cells co-expressing Flag-NDRG1 WT or S336A plasmids with GFP-Cdc42 were incubated with 30 μl of ANTI-FLAG® M2 Affinity Gel (Sigma-Aldrich, A2220) and eluted with 3X FLAG® Peptide (Sigma-Aldrich, F4799) by incubating for 2 h at 4 °C in rotation. Co-IP eluents were subjected to immunoblotting. Cells expressing Flag empty vector were used as negative control. For the pulldown assays performed in **Extended data Fig 13C**, total cell lysates (1000 μg) from NIH3T3 cells co-expressing mCherry-Cdc42 WT or T17N plasmids with Flag-NDRG1 WT were subjected to FLAG pulldowns as described above. For the identification of Flag-NDRG1 WT or S336A-interacting partners in **Extended data Fig 13A**, 700 ug of total cell lysates from NIH3T3 expressing Flag-NDRG1 WT or S336A plasmids were subjected to FLAG pulldowns as described above, and co-IP eluents were subjected to mass spectrometry. Non transfected cells were used as negative controls. For the identification of Cdc42-interacting partners in **Fig 4E**, 700 ug of total cell lysates from NIH3T3 expressing GFP-Cdc42 or eGFP empty vector alone were incubated with 25 μl of GFP-Trap Magnetic Agarose (Chromotek, gtma-20) for 2 h at 4 °C in rotation after which beads were washed and subjected to on-beads digestion for mass spectrometry.

### Sample preparation for phosphoproteomics

Liver tissue (500 μg), liver MAM fractions (700 μg) or co-IP eluents from FLAG pulldowns performed in total cell lysates (700 μg) of siCon or si*Rictor* NIH3T3 cells co-expressing Flag-NDRG1 WT were homogenized in 2% SDS + 5 mM DTT to retrieve proteins in solution supplemented with complete EDTA-free protease inhibitor (Roche, 11873580001) and phosphatase inhibitor cocktails 2 and 3 (Sigma-Aldrich, P5726 and P0044). Homogenized sample was left incubating for disulfide bond reduction for 60 min at room temperature. Proteins were alkylated using 20 mM iodoacetamide for 30 min in the dark. Protein digestion was then carried out utilizing the S-trap mini cartridges (ProtiFi) as recommended by manufacturer’s instructions. Phosphorylated peptides were enriched from the entire S-trap eluate using titanium dioxide beads (TiO_2_, GL Sciences) as previously described^37^. Following TiO_2_ enrichment, peptides were concentrated with a speed vac, desalted in HLB resin (Waters) and concentrated in a speed vac once more prior for analyzing peptides by nano liquid chromatography coupled online with tandem mass spectrometry (nLC-MS/MS).

### nLC-MS/MS acquisition

Samples were resuspended in 10 μl of water + 0.1% TFA and loaded onto a Dionex RSLC Ultimate 300 (Thermo Scientific, San Jose, CA, USA) coupled online with an Orbitrap Fusion Lumos (Thermo Scientific). Chromatographic separation was performed with a two-column system, consisting of a C18 trap cartridge (300 μm ID, 5 mm length) and a picofrit analytical column (75 μm ID, 30 cm length) packed in-house with reversed-phase Repro-Sil Pur C18-AQ 3 μm resin. Peptides were separated using a 180 min gradient from 2-28% buffer-B (buffer-A: 0.1% formic acid, buffer-B: 80% acetonitrile + 0.1% formic acid) at a flow rate of 300 nl/min. The mass spectrometer was set to acquire spectra in a data-dependent acquisition (DDA) mode. Briefly, the full MS scan was set to 300-1200 m/z in the orbitrap with a resolution of 120,000 (at 200 m/z) and an AGC target of 5×10^5^. MS/MS was performed in the ion trap using the top speed mode (2 secs), an AGC target of 10e4 and an HCD collision energy of 30. Two additional targeted scans were added in each instrument duty cycle to detect the low abundance NDRG1 S336ph peptide; a selected ion monitoring (SIM) scan for the intact mass quantification and a targeted MS/MS scan for the identification of the peptide. Raw files were uploaded in the public database Chorus (https://chorusproject.org) at the Project ID #1713.

### Phosphoproteomics data analysis

Raw files were searched using the software Proteome Discoverer v2.4 (Thermo Scientific) using standard settings for tolerances, modifications and filters, and phosphorylation on serine/threonine/tyrosine as dynamic modifications. The database used was the SwissProt mouse proteome (downloaded August 2019). Peptide abundance was obtained using the intensity of the extracted ion chromatogram; values were then log2 transformed, normalized and missing values were imputed as previously described^38^. Comparisons between groups were performed in a binary manner; each sample type (basal, corn oil, palmitic acid and refed) were compared with the fasted condition utilizing a two-tails heteroscedastic t-test (significant, if P-value < 0.05). The data distribution in each dataset was assumed to be normal but this was not formally tested.

Significantly modified proteins were selected by P<0.05 followed correction using the Benjamini-Hochberg procedure. When FDR correction led to a no hit, inspection of uncorrected P-values distribution was performed: if an anti-conservative distribution was observed, we applied alternative method of false discovery rate control by combining threshold for significance (P<0.05) with fold-change cut-off (fold-change>1.5) as previously suggested^39^. The phosphorylation state change (ΔPs) value for individual proteins was calculated as previously described^19^, as the sum of log2(fold change) value of all phosphopeptides with statistically significant changes (P<0.05) compared to control. If none of the phosphopeptide P values are below 0.05, the ΔPs value will be zero. We applied a stringent cut-off for ΔPs value at 2 standard deviations (2σ) to represent the concept of cumulative phosphorylation. Gene ontology was performed using BINGO or Enrichr^40^. In the enrichment map-based network visualization of gene ontology enrichment of differentially modulated phosphosites, blue edges show similarity between decreased phosphosites while red nodes show similarity between increased phosphosites; node size indicates the number of proteins per node; major clusters are circled, and the associated name represent the major functional association. The enrichment map was generated in Cytoscape (3.8.1) using Enrichment map plugin (3.3.0)^18^ using the following thresholds: P-value <0.05, FDR <0.001. Data handling and statistical analyses were performed using Python (Python software foundation; v.3.7.4 available at https://www.python.org/) and scientific python stack: SciPy (v.1.3.1)^41^, NumPy (v.1.17.2)^42^, and Matplotlib (v.3.1.1). Phosphosites showing significant regulation between groups were used to predict the kinase responsible for their catalysis using the iGPS software^6^. Significantly regulated phosphorylation events were used to predict the kinases responsible for their catalysis using iGPS^6^. Positive kinase scores represent most confident and frequent predictions for upregulated phosphosites, while blue for downregulated phosphosites. The higher the cumulative score retrieved from iGPS and the more intense the color coding of the bubbles in the network. Upregulated and downregulated refers to numerator and denominator as defined in the header of each panel. The bubble size is scaled based on the number of phosphorylation events predicted to be catalyzed by the given kinase. The connector lines represent previously associated genetic interactions between listed proteins retrieved from the database STRING v11^43^. The network was displayed using Cytoscape^44^.

### Biochemical analysis

Blood glucose levels were measured using Ascensia Contour glucometer (Bayer). Serum insulin (ALPCO, 80-INSNS-E01), free fatty acids (FFA; FUJIFILM, NEFA-HR (2)), serum triglyceride (TG; Sigma-Aldrich, T2449, F6428) and liver TG (BioVision, K622) were evaluated as per manufacturer’s instructions.

### Seahorse respirometry

In vitro Seahorse XF Cell Mito Stress Test (Agilent Technologies) was performed according to the manufacturer’s instructions. Briefly, 1.2 × 10^4^ NIH3T3 cells per 100 μl of growth medium were transfected with DNAs or siRNAs as described above and seeded onto a Seahorse XF96 Cell Culture Microplate (Agilent Technologies, 101085-004) for 32 h. After 16 h of stress in low glucose growth medium (1 g/L; Agilent Technologies, 103577), cells were washed once with PBS, cultured in 165 μl of XF Base Medium (Agilent Technologies, 103335) supplemented with 1 g/L D-glucose, 2 mM sodium pyruvate (Gibco, 11360) and 4 mM L-Glutamine (Gibco, 25030) and incubated at 37 °C without CO_2_ for 1 h. Immediately prior to initiating the Mito Stress Test, oleic acid (0.25 mM) was added to cells and microplate was loaded into XF analyzer. Basal oxygen consumption rates (OCR) measurements were recorded 4 times (mix: 3 min; wait: 2 min; measure: 3 min), followed by sequential injection of mitochondrial respiration modulators oligomycin (1 μM), carbonyl cyanide-4 (trifluoromethoxy) phenylhydrazone (FCCP; 20 μM), and rotenone and antimycin (1 μM) with 4 readings (mix: 3 min; wait: 2 min; measure: 3 min) after each injection. OCR values were normalized to cell number per well estimated with CyQUANT™ Cell Proliferation Assay (Invitrogen, C7026) as per manufacturer’s instructions. Parameters of mitochondrial function were calculated as per manufacturer’s instructions. OCRs of liver explants were performed as described previously^34^.

### Histological analyses

Immunohistochemistry to detect FLAG was performed using a M.O.M.® (Mouse on Mouse) ImmPRESS® HRP (Peroxidase) Polymer Kit (Vector Laboratories, MP-2400). Paraffin-embedded livers were cut into 5-μm-thick sections and subjected to deparaffination in xylene followed by rehydration in a series of graded alcohols and water. For antigen unmasking, sections were incubated in citrate-based antigen unmasking solution (pH 6.0; Vector Laboratories, H-3300) in combination with high temperature for 20 min. After blocking in BLOXALL® Endogenous Blocking Solution (Vector Laboratories, SP-6000) for 10 min followed by a 1 h-incubation in M.O.M. Mouse IgG Blocking Reagent, sections were stained with mouse monoclonal anti-DYKDDDDK Tag antibody (1:100; Cell Signaling Technology, 8146) in 2.5% Normal Horse Serum M.O.M. solution overnight at 4 °C. DYKDDDDK signal was revealed by incubation with M.O.M. ImmPRESS Reagent for 10 min and enhanced with ImmPACT® DAB EqV Peroxidase (HRP) Substrate (Vector laboratories, SK-4103) for 1 min. Sections were counterstained with hematoxylin, dehydrated, mounted with Permount™ mounting medium (Fisher, SP15) and imaged in a Zeiss Axiolab 5 microscope/Axiocam 305 color camera (Carl Zeiss Microscopy, Thornwood, NY).

### Oil red O staining

Oil red O staining was performed as described previously^45^.

### Confocal fluorescence microscopy

Fluorescence microscopy was performed as described previously^36^. For FLAG detection, DYKDDDDK Tag Rabbit antibody was used at 1:100 dilution (Cell Signaling Technology, 14793). Where indicated, 30 min prior to fixation with 4% paraformaldehyde, cells were incubated with 100 nM MitoTracker™ Red CMXRos (Invitrogen, M7512) to assess mitochondrial membrane potential. Mounted coverslips were imaged on a Leica TCS SP8 Confocal Laser Scanning Microscope (Leica Microsystems, Buffalo Grove, IL) with X63 objective and 1.4 numerical aperture. Quantification of MitoTracker™ Red CMXRos fluorescence intensity per cell was performed using ImageJ software (NIH) and expressed as mean integrated density. For detection of BODIPY FL C_16_ in vivo, sections from freshly isolated livers were mounted with Fluoromount-G medium (SouthernBiotech, 0100) and imaged on Leica TCS SP8 Confocal Laser Scanning Microscope with X10 objective and 1.4 numerical aperture.

### Live cell imaging

Cells were transfected with siRNAs and/or DNA plasmids as described above and seeded onto a glass-bottom 35mm culture dish (MatTek Corporation, P35G-1.5-14-C) for 48 h. After washing with PBS, the cells were incubated in serum-free DMEM in the presence of MitoTracker™ Green FM (500 nM; Invitrogen, M7514) or ER-Tracker™ Green (500 nM; Invitrogen, E34251) for 30 min to stain the mitochondria and endoplasmic reticulum, respectively. Just prior to visualization, the cells were washed once with PBS and incubated in red phenol-free DMEM (Gibco, 31053) supplemented with 4mM L-Glutamine and 12mM HEPES (pH=7.4), imaged using a Leica TCS SP8 Confocal Laser Scanning Microscope (Leica Microsystemsf, Buffalo Grove, IL) and single planes were acquired with ×63 objective and 1.4 numerical aperture. For experiments comprising time-lapse imaging, cells were tracked at a rate of 1 frame per 13 or 27 seconds (for single or simultaneous dual-channel acquisition, respectively) over 10 minutes.

### Image analysis

Images were analyzed in ImageJ software (NIH). Individual frames were denoised by applying Gaussian filter and a region of interest (ROI) of 35 μm^2^ was selected across the different experimental conditions. After image auto-thresholding, the quantification of mitochondrial number and mitochondrial morphology parameters was performed using “analyze particles” macro as previously described^46^. Mitochondrial elongation was calculated as the inverse of average circularity^47^. Mitochondrial fission and fusion frequency were calculated as previously described^48^ and expressed as number of events per cell per second. Percentage colocalization was calculated using the JACoP plugin as previously described^36^.

### Transmission electron microscope

Freshly isolated livers were fixed with 2% paraformaldehyde and 2.5% glutaraldehyde in 0.1 M sodium cacodylate buffer, post-fixed with freshly prepared 2% osmium tetroxide, 1.5% potassium ferrocyanide, 0.15 M sodium cacodylate, 2 mM CaCl_2_, followed by 1% thiocarbohydrazide, and then 2 % osmium tetroxide, *en bloc* stained with 1% uranyl acetate and further stained with lead aspartate. Samples were dehydrated through graded series of ethanol and embedded in LX112 resin (LADD Research Industries, Burlington VT). Ultrathin (55 nm) sections were cut on a Leica ARTOS 3D ultramicrotome and collected onto silicon wafers. Sections were examined on Zeiss Supra 40 Field Emission Scanning Electron Microscope (Carl Zeiss Microscopy, LLC North America) in backscatter mode using an accelerating voltage of 8.0 KV. The assessment of mitochondrial number and morphology was performed using ImageJ software (NIH). The number of mitochondria were counted manually in a ROI of 71.2 μm^2^ at a magnification of 2.5 kx. Quantitative analysis of mitochondria shape descriptors was performed by manual tracing of individual mitochondria using the freehand tool at a magnification of 5 kx. Contact sites between mitochondria and ER (defined to be at 10-30 nm distance^49^) were quantified, normalized to the total number of mitochondria counted and expressed as %. For 3D reconstruction, regions of interest were collected using ATLAS 5.0, with a pixel size of 6.0 and dwell time of 6 μs. Stacks were aligned and segmentation was done using IMOD^50^. Tomographic reconstruction was performed as described^51^.

**Lipidomic analyses** Lipid extracts from whole liver homogenates, MAM, Mp and ER fractions were prepared using modified Bligh and Dyer method, spiked with appropriate internal standards, and analyzed on a platform comprising Agilent 1260 Infinity HPLC integrated to Agilent 6490A QQQ mass spectrometer controlled by Masshunter v 7.0 (Agilent Technologies, Santa Clara, CA). Glycerophospholipids and sphingolipids were separated with normal-phase HPLC as described before (Chan et al, 2012), with a few modifications. An Agilent Zorbax Rx-Sil column (2.1 × 100 mm, 1.8 µm) maintained at 25°C was used under the following conditions: mobile phase A (chloroform: methanol: ammonium hydroxide, 89.9:10:0.1, v/v) and mobile phase B (chloroform: methanol: water: ammonium hydroxide, 55:39:5.9:0.1, v/v); 95% A for 2 min, decreased linearly to 30% A over 18 min and further decreased to 25% A over 3 min, before returning to 95% over 2 min and held for 6 min. Separation of sterols and glycerolipids was carried out on a reverse phase Agilent Zorbax Eclipse XDB-C18 column (4.6 × 100 mm, 3.5um) using an isocratic mobile phase, chloroform, methanol, 0.1 M ammonium acetate (25:25:1) at a flow rate of 300 μl/min.

### Quantification of lipid species

was accomplished using multiple reaction monitoring (MRM) transitions (Chan et al, 2012; Hsu et al, 2004; Guan et al, 2007) under both positive and negative ionization modes in conjunction with referencing of appropriate internal standards: PA 14:0/14:0, PC 14:0/14:0, PE 14:0/14:0, PG 15:0/15:0, PI 17:0/20:4, PS 14:0/14:0, BMP 14:0/14:0, APG 14:0/14:0, LPC 17:0, LPE 14:0, LPI 13:0, Cer d18:1/17:0, SM d18:1/12:0, dhSM d18:0/12:0, GalCer d18:1/12:0, GluCer d18:1/12:0, LacCer d18:1/12:0, D7-cholesterol, CE 17:0, MG 17:0, 4ME 16:0 diether DG, D5-TG 16:0/18:0/16:0 (Avanti Polar Lipids, Alabaster, AL). Lipid levels for each sample were calculated by summing up the total number of moles of all lipid species measured by all three LC-MS methodologies, and then normalizing that total to mol %. The final data are presented as mean mol % with error bars showing mean ± SEM.

### Illustration

The proposed model in Fig. 4h was created with BioRender.com.

### Statistics

All data are mean values and are from a minimum of three independent experiments unless otherwise stated. Statistical significance was assessed by two-tailed unpaired Student’s t test, one-way or two-way ANOVAs followed by Tukey’s, Šídák’s or Dunnett’s multiple-comparison test. *n* numbers indicate biological replicates. **Table S2**.

