## Supplemental Figures for "mTORC2 couples fasting to mitochondrial fission"

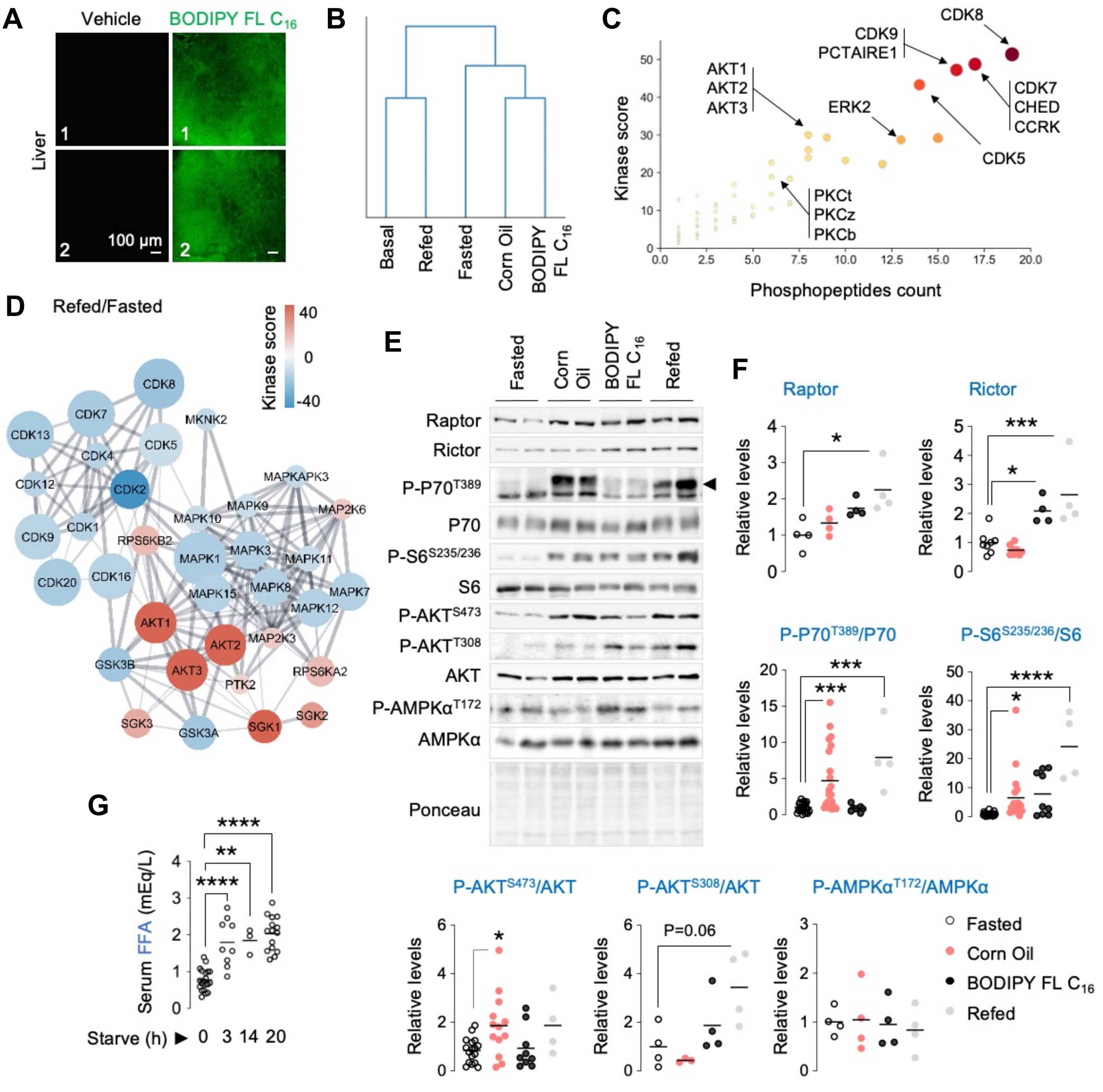


**Extended Data Fig. 1 | Kinases responsive to fasting or fatty acids. (A)** Direct BODIPY fluorescence in livers of 8 mo-old C57BL/6 male mice subjected to 30 min gavage with vehicle or 10 mg/kg of BODIPY FL $C_{16}$ after 14-16 h fasting (n=5). **(B)** Dendrogram of the hierarchical clustering analysis (Euclidean distance) between experimental groups performed on the phosphorylation status of 863 phosphosites. **C)** iGPS prediction of upstream kinases performed at the individual kinase level (as opposed to Main Fig. 1d panel, which represents analyses at the level of kinase families). **(D)** Pairwise comparisons showing upregulated or downregulated kinase networks in mice refed high fat diet (HFD) for 30 min after 14-16 h fasting. **(E)** Representative IB, and **(F)** quantifications for indicated proteins in livers of 2-8 mo-old C57BL/6 male or female mice kept fasted for 14-16 h or treated with corn oil or BODIPY FL $C_{16}$ or refed a HFD for 30 min after fasting (n=4-23). Ponceau is the loading control. **(G)** Circulating free fatty acids (FFA) in 2-7 mo-old male mice fed or fasted for indicated periods (n=3-20 mice). Individual replicates and mean values are indicated. *P<0.05, **P<0.01, ***P<0.001 and ****P<0.0001. One-way ANOVA followed by Šídák's multiple comparisons test **(F)**. One-way ANOVA followed by Tukey's multiple comparisons test **(G)**. Please refer to Table S2_statistical summary. Please refer to excel files, proteomics1b.

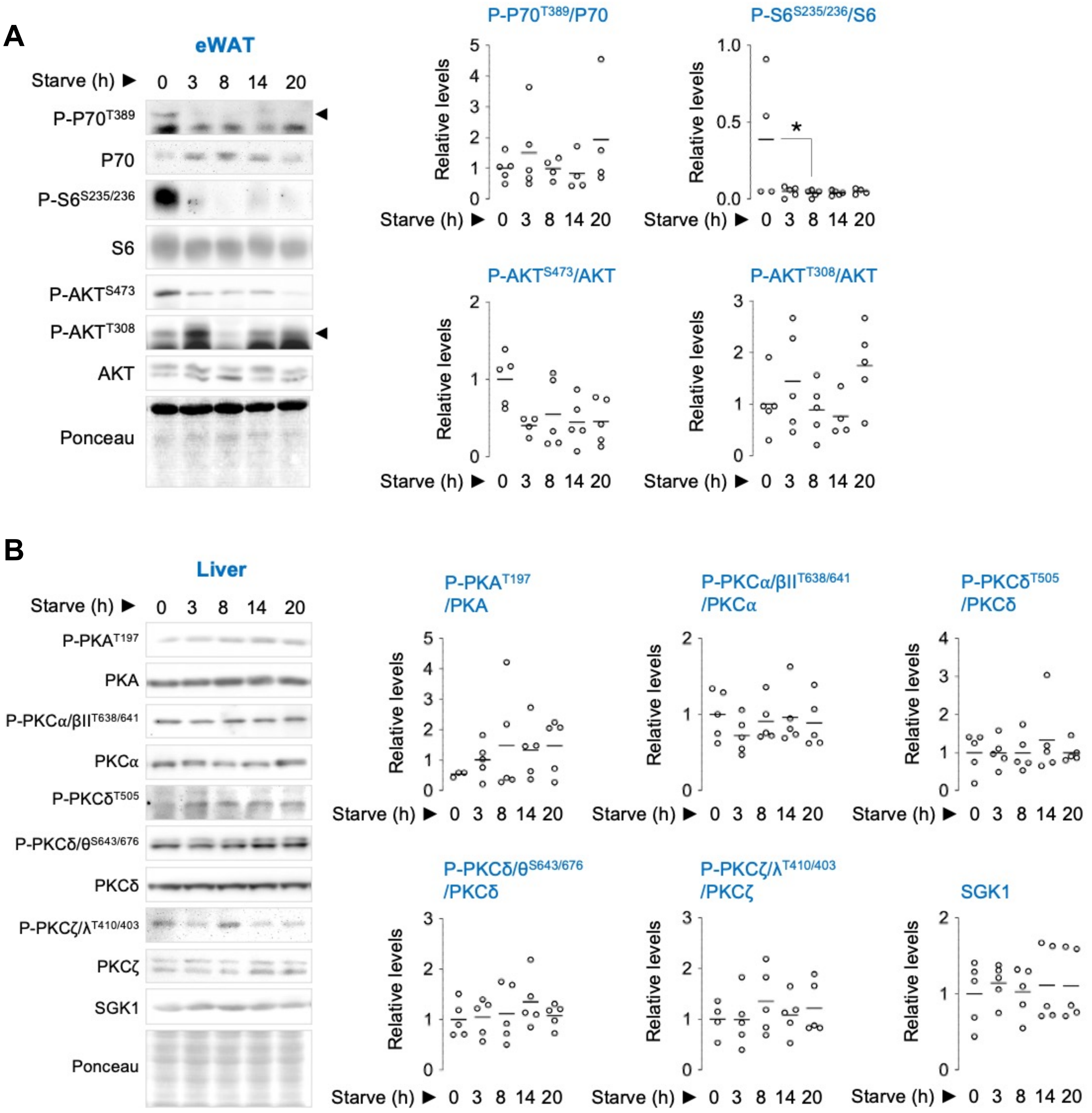


**Extended Data Fig. 2 | Protein kinases A and Cs are not activated by fasting in liver, while mTORC1/C2 do not respond to fasting in eWAT. (A)** IB and quantification for indicated markers for mTORC1 and mTORC2 signaling in epididymal white adipose tissue (eWAT) of 5-10 mo-old male mice that were fed or fasted for indicated periods (n=4-5 mice). **(B)** IB and quantification for the indicated proteins in liver of 2-10 mo-old mice fed or fasted for the indicated durations (n=4-27 mice). Ponceau is loading control. Individual replicates and means are shown. *P<0.05, One-way ANOVA followed by Tukey's multiple comparisons test. Please refer to Table S2_statistical summary.

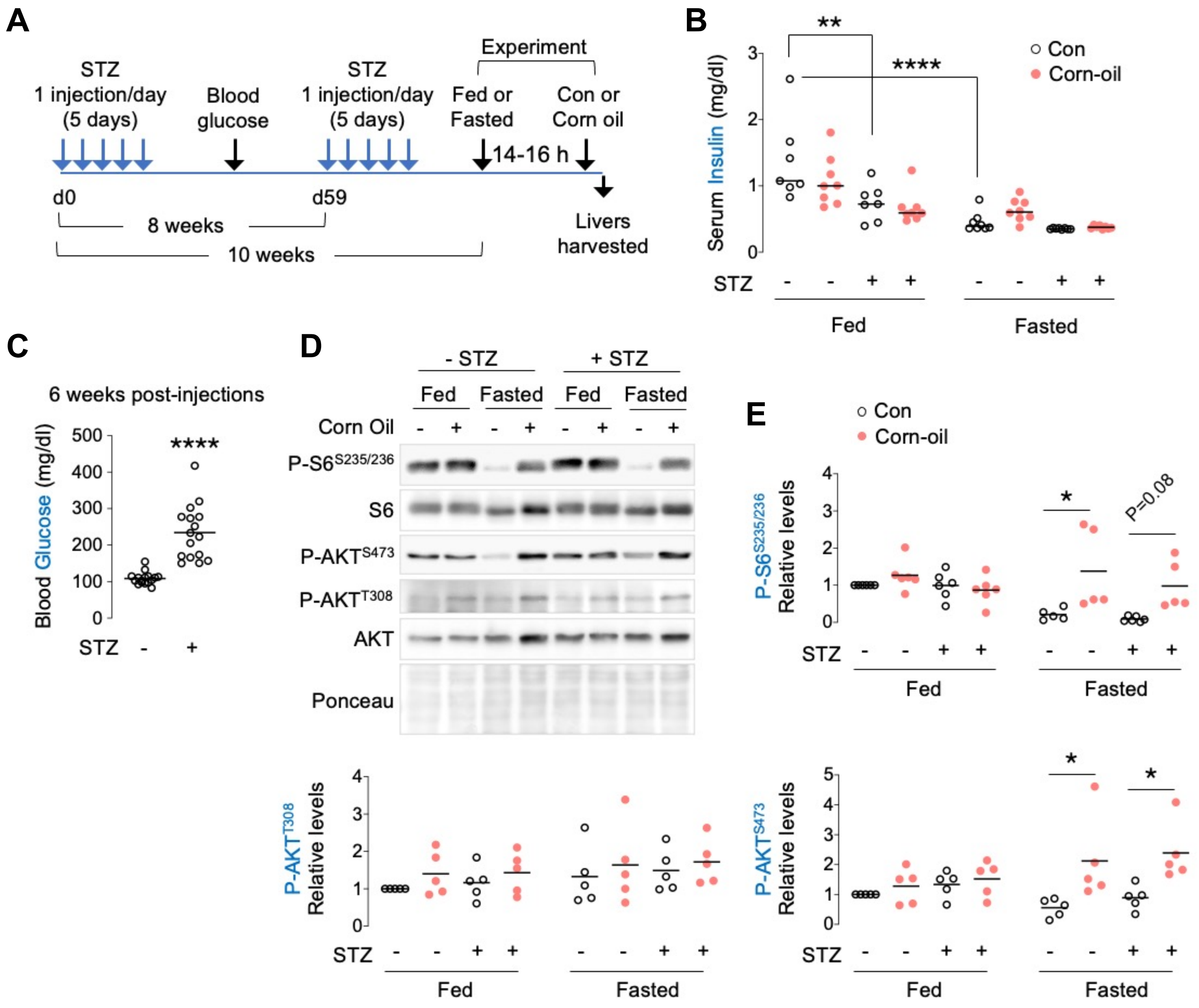


**Extended Data Fig. 3 | mTORC1/C2 signaling in presence of lipid appears to be insulin-independent. (A)** Experimental plan for generation of insulin-deficient diabetic mice using streptozotocin (STZ). **(B)** Serum insulin levels in 3 mo-old STZ-treated C57BL/6 male mice that were fed or fasted for 14-16 h and then gavaged with corn oil for 30 min (n=7-8). **(C)** Blood glucose levels 6 weeks after the first injection with STZ (n=16). **(D)** Representative IB, and **(E)** quantification for indicated proteins normalized to corresponding total protein in livers of 3 mo-old STZ-treated C57BL/6 male mice that were fed or fasted for 14-16 h and then gavaged with corn oil for 30 min (n=5-6). Ponceau is the loading control. Individual replicates and mean values are indicated. *P<0.05, **P<0.01 and ****P<0.0001, 2-way ANOVA followed by Tukey's correction **(B and E)**; unpaired Student's t-test **(C)**. Please refer to Table S2_statistical summary.

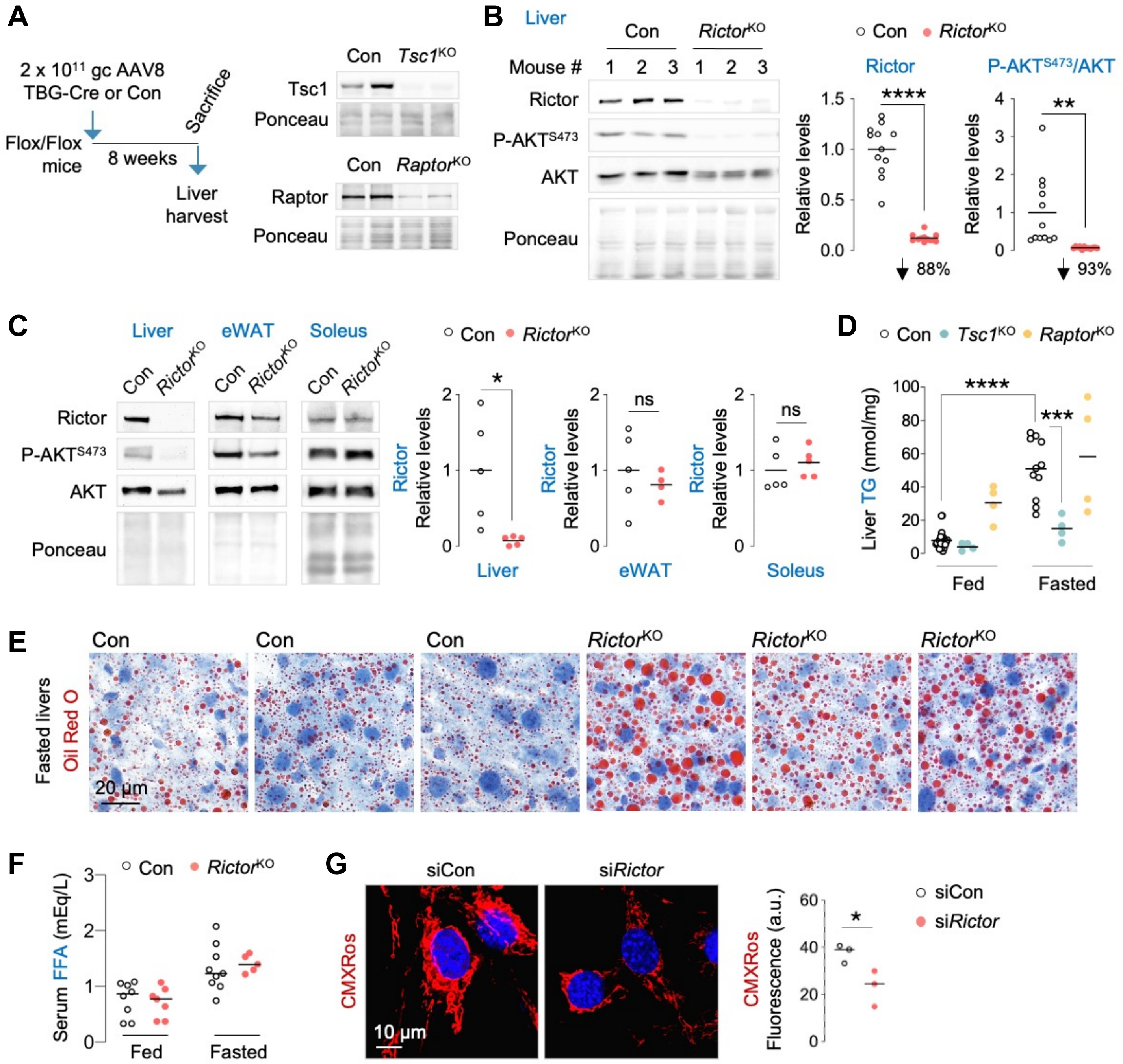


**Extended Data Fig. 4 | Impact of loss of mTORC2 signaling on lipid metabolism and mitochondrial membrane potential. (A-B)** Generation of liver-specific knock-out (KO) of *Tsc1*, *Raptor* or *Rictor*. Representative IB to validate deletion of indicated genes in livers of 4-6 mo-old **(A)** Con, *Tsc1*$^{KO}$, *Raptor*$^{KO}$ or **(B)** *Rictor*$^{KO}$ male and female mice. Quantification and % reduction of protein levels for Rictor and P-AKT$^{S473}$/AKT in *Rictor*$^{KO}$ livers is shown in **B** (n=12 mice). **(C)** IB for mTORC2 signaling and quantification of Rictor protein levels in liver, eWAT and soleus of 4-6 mo-old Con and *Rictor*$^{KO}$ male mice fasted for 14-16 h (n=4-5 mice). **(D)** Liver TGs in 3-7 mo-old Con, *Tsc1*$^{KO}$ and *Raptor*$^{KO}$ male and female mice that were fed or fasted for 14-16 h (n=4-24). **(E)** Representative immunohistochemical staining of Oil red O in livers of 6 mo-old Con or *Rictor*$^{KO}$ male mice fasted for 14-16 h (n=5 mice). Three distinct Con and *Rictor*$^{KO}$ livers are shown. **(F)** Serum FFA levels in fed or 14-16 h fasted 3-6 mo-old Con or *Rictor*$^{KO}$ male mice (n=5-9). **(G)** MitoTracker CMXRos fluorescence in serum-starved and oleic-acid treated siControl or si*Rictor* NIH3T3 cells (58-60 cells from 3 independent experiments). Ponceau is loading control. Individual replicates and means are shown. *P<0.05, **P<0.01, ***P<0.001 and ****P<0.0001. Unpaired Student's t-test **(B, C and G)**; 2-way ANOVA followed by Tukey's multiple comparison test **(D).** ns = not significant. Please refer to Table S2_statistical summary.

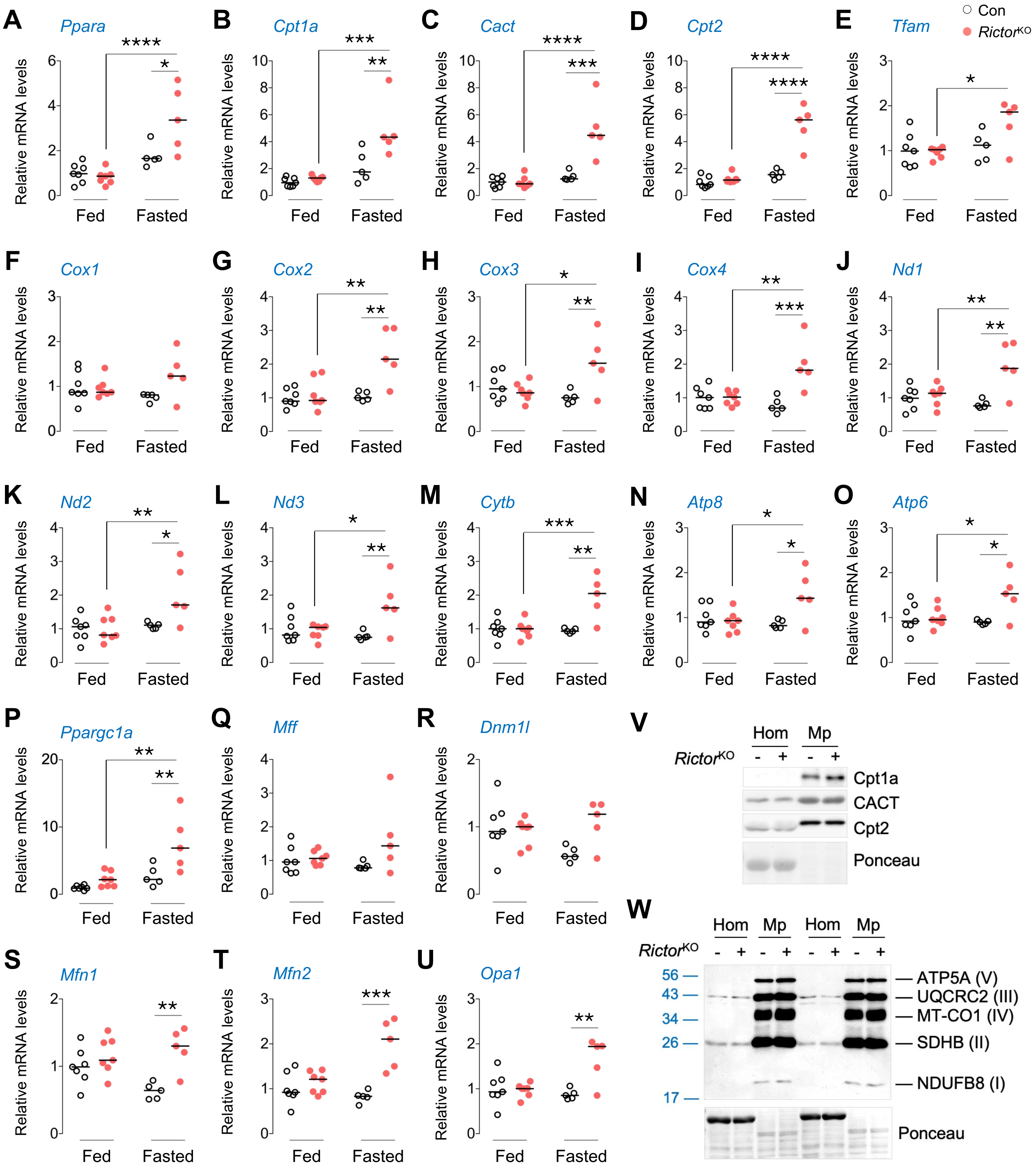


**Extended Data Fig. 5 | Effect of loss of mTORC2 on expression of genes and proteins related to mitochondrial oxidative metabolism. (A-U)** Relative mRNA expression of indicated genes in livers of 4-6 mo-old Con and *Rictor*$^{KO}$ male mice that were fed or fasted for 14-16 h (n=5-7). **(V)** Representative IB for proteins involved in mitochondrial fatty acid uptake, and **(W)** OXPHOS in whole homogenates (Hom) and pure mitochondrial (Mp) fractions from livers of 5-6 mo-old Con and *Rictor*$^{KO}$ male mice after 14-16 h fasting (n=3-5). Ponceau is the loading control. Individual replicates and mean values are shown. *P<0.05, **P<0.01, ***P<0.001 and ****P<0.0001. 2-way ANOVA followed by Tukey's multiple comparison **(A-U)**. Please refer to Table S2_statistical summary.

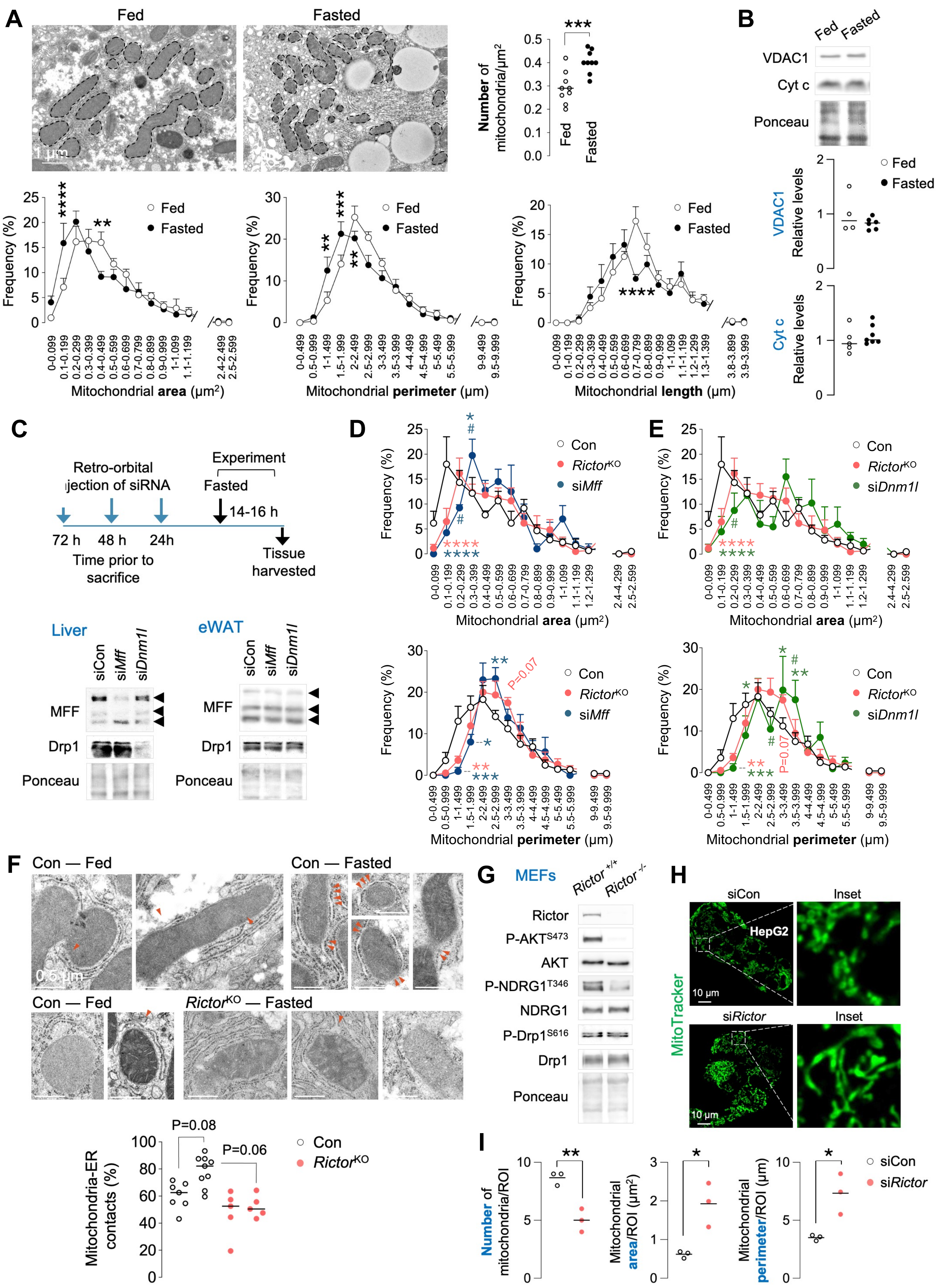
A
Fed
Fasted
1 µm
Number of mitochondria/µm²
Fed
Fasted
Frequency (%)
Mitochondrial area (µm²)
Mitochondrial perimeter (µm)
Mitochondrial length (µm)
B
VDAC1
Cyt c
Ponceau
VDAC1 Relative levels
Cyt c Relative levels
C
Retro-orbital injection of siRNA
Experiment
Fasted
14-16 h
72 h
48 h
24h
Time prior to sacrifice
Tissue harvested
Liver
eWAT
siCon
siMff
siDnm1l
MFF
Drp1
Ponceau
D
E
Con
RictorKO
siMff
siDnm1l
P=0.07
F
Con — Fed
Con — Fasted
RictorKO — Fasted
0.5 µm
Mitochondria-ER contacts (%)
P=0.08
P=0.06
G
MEFs
Rictor +/+
Rictor -/-
Rictor
P-AKTS473
AKT
P-NDRG1T346
NDRG1
P-Drp1S616
Drp1
Ponceau
H
siCon
Inset
HepG2
siRictor
MitoTracker
10 µm
I
Number of mitochondria/ROI
Mitochondrial area/ROI (µm²)
Mitochondrial perimeter/ROI (µm)
siCon
siRictor

**Extended Data Fig. 6 | Fasting associates with mitochondrial morphology reflecting increased fission, which occurs in an mTORC2-dependent manner. (A)** Liver EMs of fed or 14-16 h fasted 4-7-mo-old male mice (n=9). Quantification for mitochondrial number is shown. Frequency histograms depicting distribution of mitochondrial area, perimeter and length. Mean ± SEM is shown. **(B)** Representative IB and quantification for indicated mitochondrial markers in livers of fed or 14-16 h fasted mice (n=4-7). **(C-E) (C)** Experimental plan and representative IB for indicated proteins in livers of 4-9 mo-old male mice injected with siCon, si*Mff* or si*Dnm1l* and fasted for 14-16 h (n=3-4). Frequency histograms depicting distribution of mitochondrial area and perimeter in Con, *Rictor*$^{KO}$ and **(D)** si*Mff* or **(E)** si*Dnm1l* livers. Mean ± SEM is shown. **(F)** EMs in fed or 14-16 h fasted Con and *Rictor*$^{KO}$ livers of 4-9 mo-old male mice (n=4-9 mice). Quantification for % of mitochondria-ER contacts is shown. Red arrowheads depict contact sites. **(G)** IB for indicated proteins in *Rictor* $^{+/+}$ and *Rictor* $^{-/-}$ MEFs. **(H)** Representative confocal images of siCon or si*Rictor* HepG2 cells cultured in serum-free medium for 30 min in presence of MitoTracker green. Magnified insets are shown. Quantification for mitochondrial number and mitochondria morphology parameters is shown in **(I)** (36-43 cells from 3 independent experiments). Ponceau is loading control. Individual replicates and means are shown. *P<0.05, **P<0.01, ***P<0.001 and ****P<0.0001. Unpaired student's t-test **(A, number of mitochondria** and **I)**. *P<0.05, **P<0.01, ***P<0.001 and ****P<0.0001 are versus Con. $^{\#}$P<0.05 versus *Rictor*$^{KO}$. One-way ANOVA followed by Tukey's multiple comparisons test **(D** and **E)**; 2-way ANOVA followed by Šídák's multiple comparison test **(A, histograms)**. 2-way ANOVA followed by Tukey's multiple comparison test **(F)**. Please refer to Table S2_statistical summary.

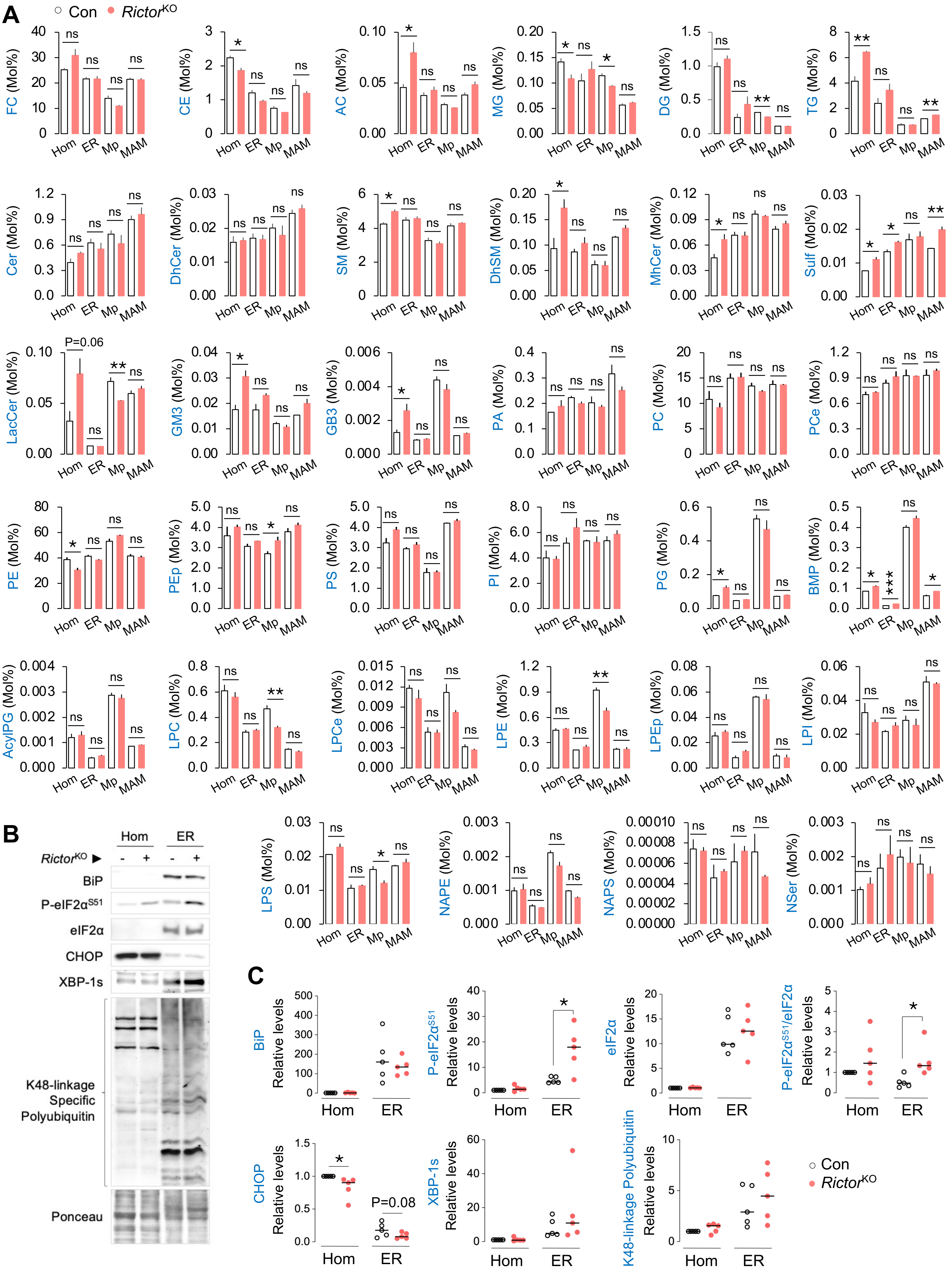
A
Con
Rictor^KO
FC (Mol%)
CE (Mol%)
AC (Mol%)
MG (Mol%)
DG (Mol%)
TG (Mol%)
Cer (Mol%)
DhCer (Mol%)
SM (Mol%)
DhSM (Mol%)
MhCer (Mol%)
Sulf (Mol%)
LacCer (Mol%)
P=0.06
GM3 (Mol%)
GB3 (Mol%)
PA (Mol%)
PC (Mol%)
PCe (Mol%)
PE (Mol%)
PEp (Mol%)
PS (Mol%)
PI (Mol%)
PG (Mol%)
BMP (Mol%)
AcylPG (Mol%)
LPC (Mol%)
LPCe (Mol%)
LPE (Mol%)
LPEp (Mol%)
LPI (Mol%)
LPS (Mol%)
NAPE (Mol%)
NAPS (Mol%)
NSer (Mol%)
Hom
ER
Mp
MAM
ns
*
**
***
B
Hom
ER
Rictor^KO ▶
- + - +
BiP
P-eIF2α^S51
eIF2α
CHOP
XBP-1s
K48-linkage Specific Polyubiquitin
Ponceau
C
BiP
P-eIF2α^S51
eIF2α
P-eIF2α^S51/eIF2α
CHOP
XBP-1s
K48-linkage Polyubiquitin
Relative levels
P=0.08
Hom
ER
Con
Rictor^KO

**Extended Data Fig. 7 | No evidence for ER stress, proteostasis failure or altered lipid metabolism in ER, MAMs or mitochondria isolated from *Rictor*$^{KO}$ livers. (A)** Lipidomics for indicated lipid species in Hom, ER, Mp and and MAM fractions from livers of 4-9 mo-old Con or *Rictor*$^{KO}$ male mice fasted for 14-16 h (n=3). Two livers were pooled to generate 1 sample. **(B)** IB and **(C)** quantification for indicated ER-stress and proteostasis markers in Hom and ER fractions from livers of 6 mo-old Con and *Rictor*$^{KO}$ mice fasted for 14-16 h (n=5 mice). Ponceau is loading control. Individual replicates and means are shown. *P<0.05, **P<0.01 and ***P<0.001. Unpaired Student's t-test **(A** and **C)**. ns = not significant. Please refer to Table S2_statistical summary.

**AC**, Acylcarnitine; **AcylPG**, Acylphosphatidyl glycerol; **BMP**, Bis[monoacylglycero]phosphate; **CE**, Cholesterol ester; **Cer**, Ceramides; **DG**, Diglyceride; **DhCer**, Dihydroceramide; **DhSM**, Dihydrosphingomyelin; **FC**, Free cholesterol; **GB3**, Globotriaosylceramide; **GM3**, Monosialodihexosylganglioside; **LacCer**, Lactosylceramide; **LPC**, Lysophosphatidylcholine; **LPCe**, Ether lysophosphatidylcholine, **LPE**, Lysophosphatidylethanolamine; **LPEp**, Plasmalogen lysophosphatidylethanolamine; **LPI**, Lysophosphatidylinositol; **LPS**, Lysophosphatidylserine; **MG**, Monoglyceride; **MhCer**, Mono hexosylceramides; **NAPE**, N-acylphosphatidylethanolamine; **NAPS**, N-acylphosphatidylserine; **NSer**, N-acylphosphatidylserine; **PA**, Phosphatidic acid; **PC**, Phosphatidylcholine; **PCe**, Ether phosphatidylcholine; **PE**, Phosphatidylethanolamine; **PEp**, Plasmalogen phosphatidylethanolamine; **PG**, Phosphatidylgylcerol; **PI**, Phosphatidylinositol; **PS**, Phosphatidylserine; **SM**, Sphingomyelin; **Sulf**, Sulfatide; **TG**, Triglycerol.

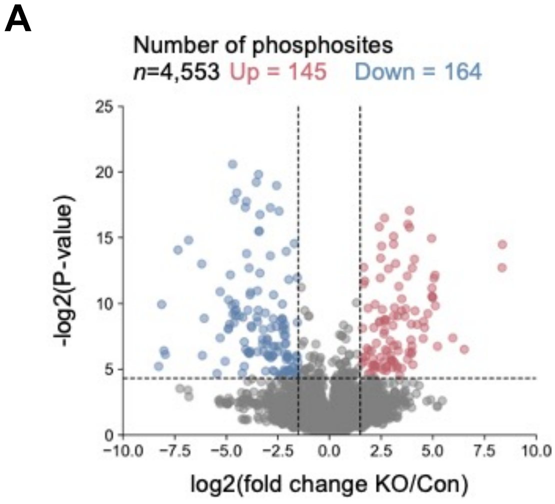


**B** Liver BCL2/adenovirus E1B 19kDa protein-interacting protein (BNIP3): -log2 transformed P-value

| Phospho-site | Fold-change KO/Con | P-value | Motif |
|---|---|---|---|
| T66 | -0.38 | 0.49 | SHCDSPPRSQTPQDTNRAEID |
| S77 | -3.54 | 2.25 | PQDTNRAEIDSHSFGEKNSTL |
| S79 | -1.57 | **5.93** | DTNRAEIDSHSFGEKNSTLSE |
| S85 | -3.96 | 2.28 | IDSHSFGEKNSTLSEEDYIER |
| T86 | 2.18 | 1.16 | DSHSFGEKNSTLSEEDYIERR |
| S88 | -6.07 | **8.87** | HSFGEKNSTLSEEDYIERRRE |

**C** Liver N-Myc Downstream Regulated Gene 1 (NDRG1): -log2 transformed P-value

| Phospho-site | Fold-change KO/Con | P-value | Motif |
|---|---|---|---|
| T328 | -0.87 | 1.81 | ASMTRLMRSRTASGSSVTSLE |
| S330 | -0.80 | 1.77 | MTRLMRSRTASGSSVTSLEGT |
| S332 | -1.30 | 1.62 | RLMRSRTASGSSVTSLEGTRS |
| S333 | 0.11 | 0.09 | LMRSRTASGSSVTSLEGTRSR |
| S336 | -0.91 | 1.43 | SRTASGSSVTSLEGTRSRSHT |

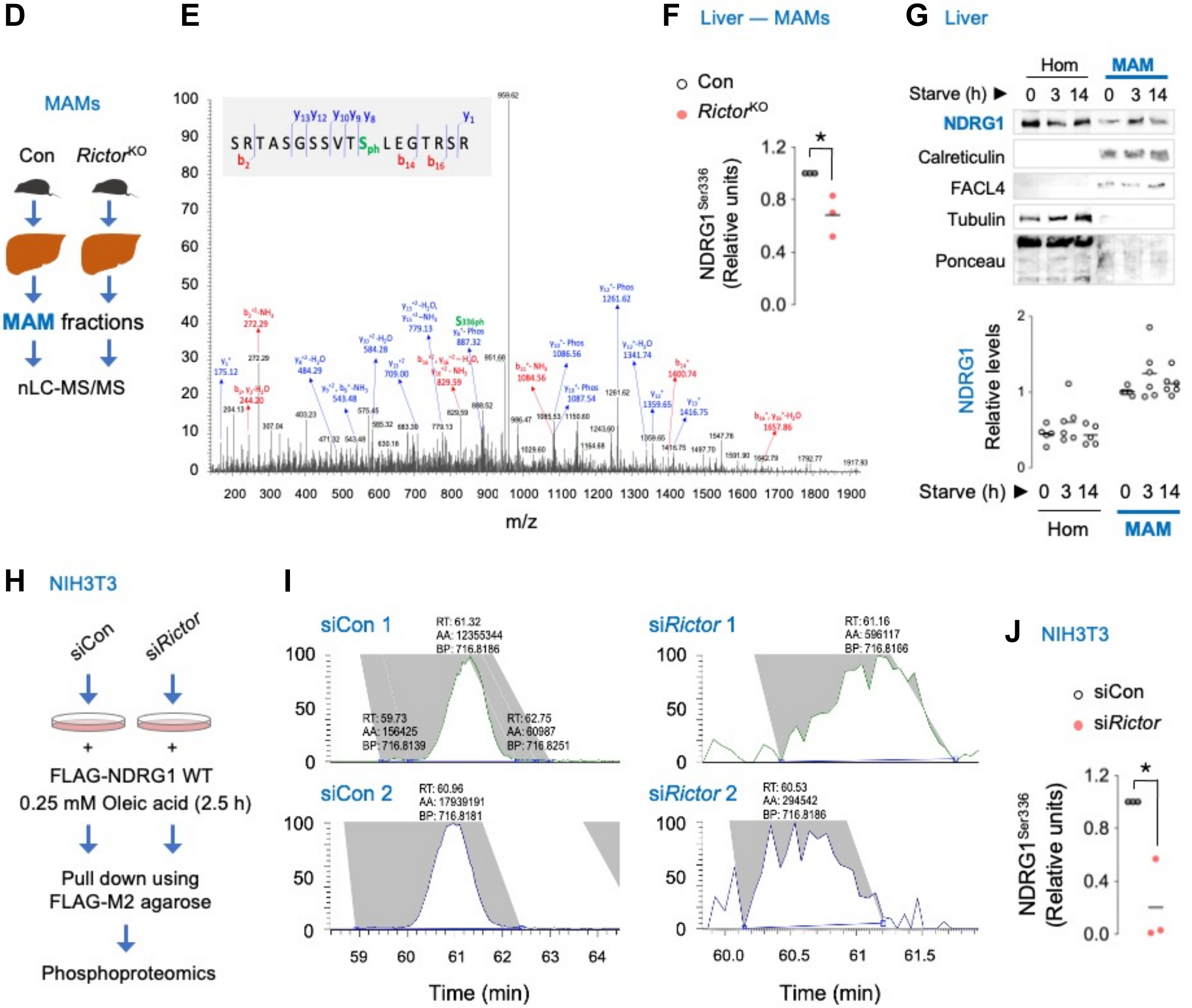

**Extended Data Fig. 8 | Phosphoproteomics reveal mTORC2 phosphorylation of NDRG1 at serine 336 in MAMs. (A)** Volcano plot for phosphoproteomics in 14-16 h fasted Con and *Rictor*$^{KO}$ livers. The numbers indicate total phosphosites. Red and blue dots represent significantly increased (P<0.05 and log2(fold change)>1.5) and decreased (P<0.05 and log2(fold change)<1.5) phosphosites, respectively (n=3). **(B, C)** Tables showing fold-change and log2 transformed P-values for indicated phosphorylations on liver **(B)** BNIP3, and **(C)** NDRG1 from 3-4-mo-old liver-specific *Rictor*$^{KO}$ mice fasted for 14-16 h. **(D)** MAMs of 4-6 mo-old Con and *Rictor*$^{KO}$ livers from 14-16 h fasted mice were subjected to phosphoproteomics. **(E)** Annotated MS/MS spectrum of phosphopeptide SRTASGSSVT**S(ph)**LEGTRSR in NDRG1, wherein **S(ph)** represents S336. **(F)** Quantification of SRTASGSSVT**S(ph)**LEGTRSR peptide from NDRG1 in MAMs are shown, with relative abundance of SRTASGSSVT**S(ph)**LEGTRSR in Con or *Rictor*$^{KO}$ MAMs (n=3 samples wherein 2 livers were pooled to generate 1 sample). **(G)** Representative IB and quantification for indicated proteins in Hom and MAM fractions from livers of 2-10 mo-old male mice fed or fasted for indicated time points. Ponceau is the loading control (n=5-6). **(H)** Experimental plan to pulldown FLAG-NDRG1 WT in siCon or si*Rictor* NIH3T3 cells co-transfected with FLAG-NDRG1 WT plasmid for assessment of phosphorylation of FLAG-NDRG1 WT via phosphoproteomics. FLAG-NDRG1 WT pulled-down from total lysates of serum-starved siCon or si*Rictor* NIH3T3 cells in presence of oleic acid for 2.5 h. **(I)** Representative extracted ion chromatograms of SRTASGSSVT**S(ph)**LEGTRSR (wherein **S(ph)** represents S336) in FLAG-NDRG1 in siCon and si*Rictor* NIH3T3 cells, and **(J)** quantification for relative abundance of SRTASGSSVT**S(ph)**LEGTRSR peptide from pulled-down FLAG-NDRG1 from siCon and si*Rictor* cells expressing FLAG-NDRG1 WT (n=3). Individual replicates and means are shown. *P<0.05, unpaired Student's t-test **(F** and **J).** Please refer to Table S2_statistical summary. Please refer to excel files, Proteomics2_Liver_PhosphoSites **(A-C),** Proteomics3_Liver_MAMs **(E** and **F),** and Proteomics4_RictorKD_Cells **(I** and **J).**

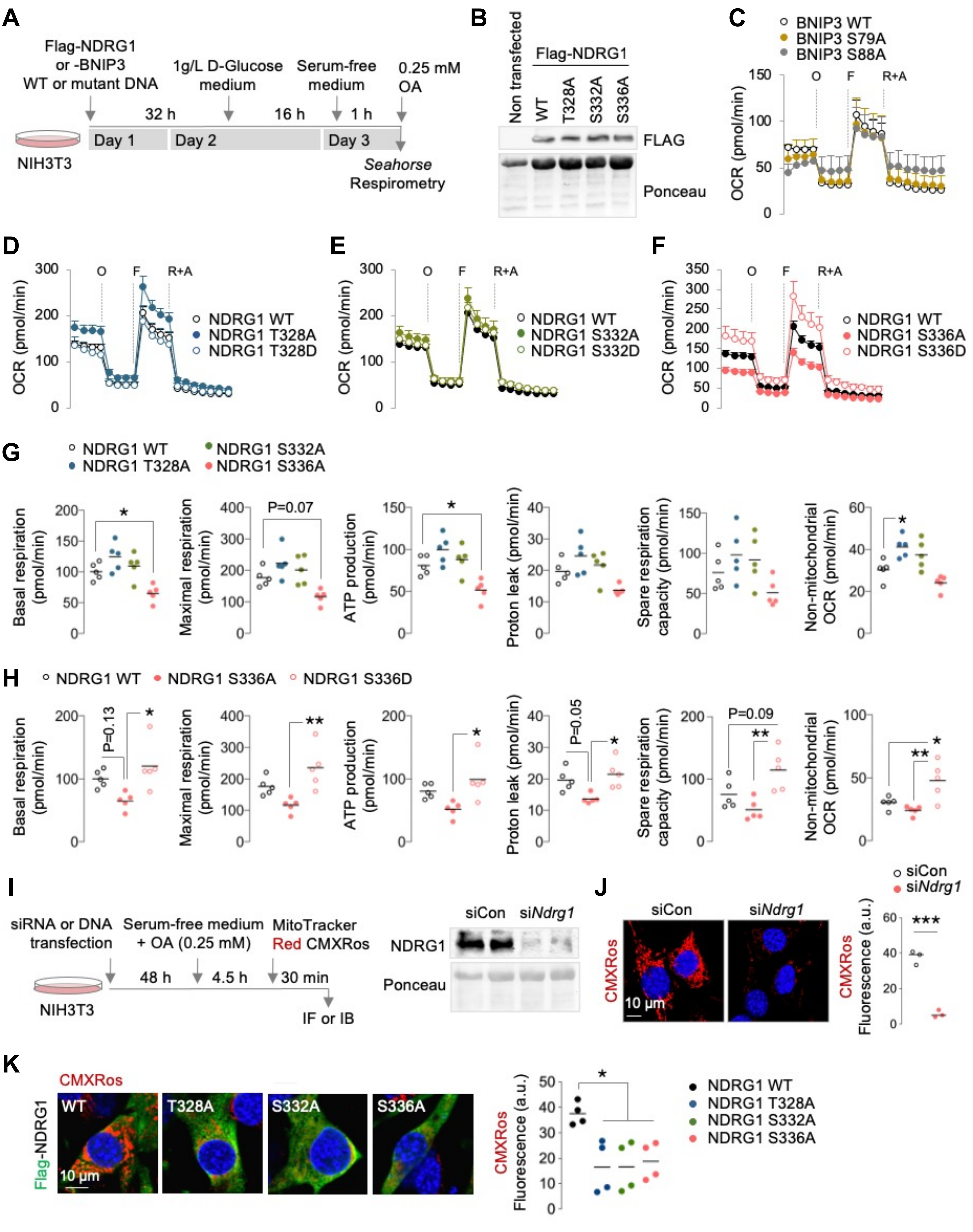

A
Flag-NDRG1 or -BNIP3 WT or mutant DNA
1g/L D-Glucose medium
Serum-free medium
0.25 mM OA
32 h
16 h
1 h
Day 1
Day 2
Day 3
NIH3T3
Seahorse Respirometry
B
Non transfected
Flag-NDRG1
WT
T328A
S332A
S336A
FLAG
Ponceau
C
BNIP3 WT
BNIP3 S79A
BNIP3 S88A
OCR (pmol/min)
O
F
R+A
D
NDRG1 WT
NDRG1 T328A
NDRG1 T328D
E
NDRG1 WT
NDRG1 S332A
NDRG1 S332D
F
NDRG1 WT
NDRG1 S336A
NDRG1 S336D
G
NDRG1 WT
NDRG1 T328A
NDRG1 S332A
NDRG1 S336A
Basal respiration (pmol/min)
Maximal respiration (pmol/min)
P=0.07
ATP production (pmol/min)
Proton leak (pmol/min)
Spare respiration capacity (pmol/min)
Non-mitochondrial OCR (pmol/min)
H
NDRG1 WT
NDRG1 S336A
NDRG1 S336D
P=0.13
P=0.05
P=0.09
I
siRNA or DNA transfection
Serum-free medium + OA (0.25 mM)
MitoTracker Red CMXRos
48 h
4.5 h
30 min
NIH3T3
IF or IB
siCon
siNdrg1
NDRG1
Ponceau
J
siCon
siNdrg1
CMXRos
10 µm
CMXRos Fluorescence (a.u.)
K
CMXRos
Flag-NDRG1
WT
T328A
S332A
S336A
10 µm
CMXRos Fluorescence (a.u.)
NDRG1 WT
NDRG1 T328A
NDRG1 S332A
NDRG1 S336A

**Extended Data Fig. 9 | Phosphorylation of NDRG1 at S336 supports mitochondrial respiration and membrane potential. (A)** Experimental plan for Seahorse mitochondrial stress tests shown through **C-H**. **(B)** IB for FLAG, and **(C)** Seahorse mitochondrial stress test in serum-starved NIH3T3 cells expressing Flag-tagged BNIP3 WT or Flag-tagged S79A or S88A mutant BNIP3 in presence of 0.25 mM oleic acid (OA), followed by sequential addition of oligomycin (O), FCCP (F), and rotenone + antimycin (R+A) to assess mitochondrial respiratory function (n=5). **(D-F)** Seahorse mitochondrial stress tests in NIH3T3 cells expressing the indicated WT or S/T>A or S/T>D mutant of NDRG1. Quantifications for mitochondrial respiratory function in cells expressing **(G)** NDRG1 WT or S/T>A mutants of NDRG1, or **(H)** NDRG1 WT or mutants NDRG1 S336A and S336D are shown (n=5). **(I)** Cartoon depicting experimental plan for confocal microscopy performed in **J** and **K**. **(J)** MitoTracker CMXRos red fluorescence in siCon or si*Ndrg1* cells (58-61 cells from 3 independent experiments). Representative blots for NDRG1 in siCon and si*Ndrg1* NIH3T3 cells are shown in **I, right**. **(K)** MitoTracker CMXRos red fluorescence in FLAG (green fluorescence)-tagged WT or phosphorylation-deficient mutants of NDRG1 (33-37 cells from 4 independent experiments). Ponceau is loading control. Individual replicates and means are shown. *P<0.05, **P<0.01 and ***P<0.001. One-way ANOVA followed by Tukey's multiple comparison test **(G, H** and **K)**; unpaired Student's t-test **(J)**. Please refer to Table S2_statistical summary.

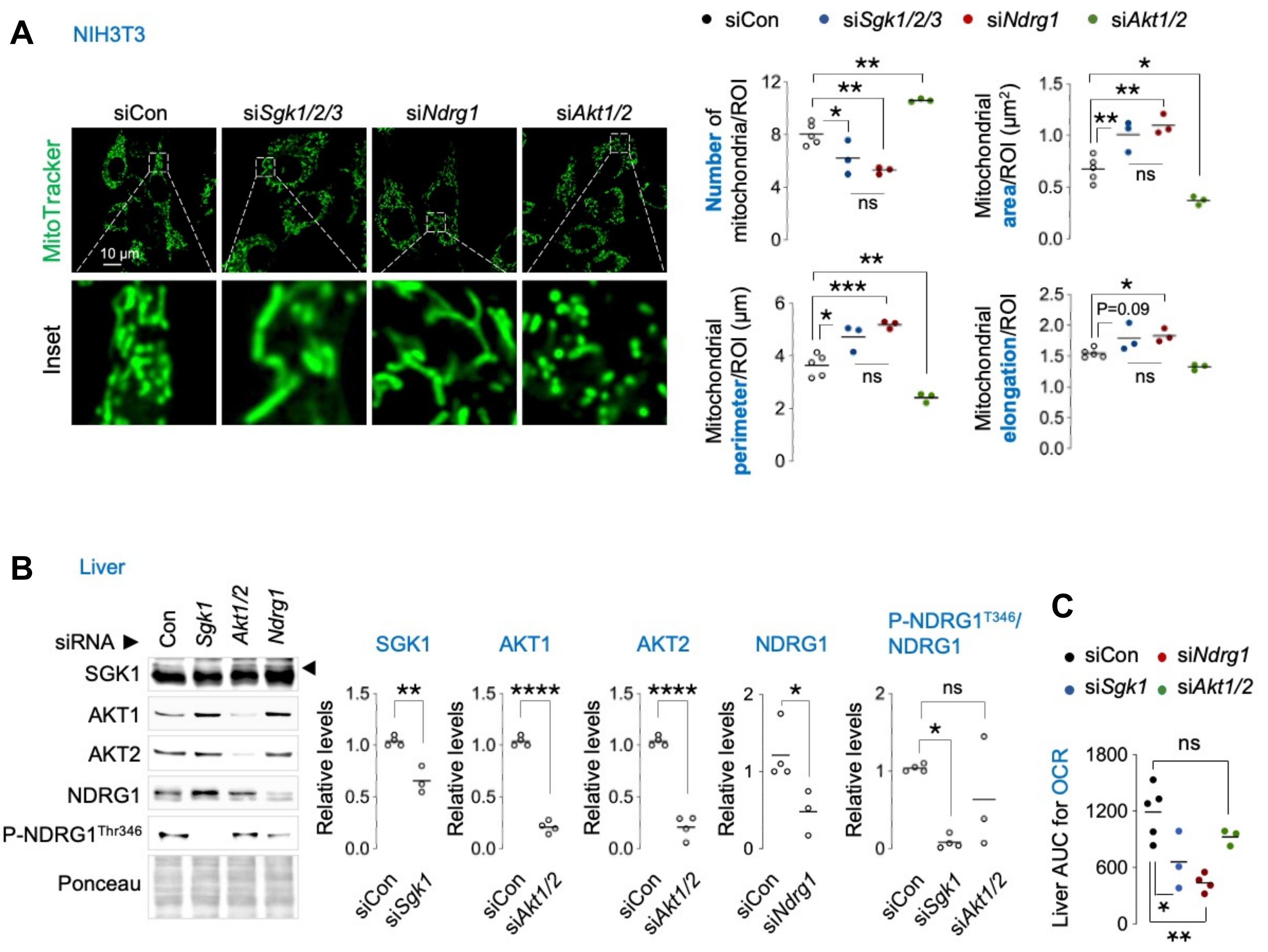


**Extended Data Fig. 10 | NDRG1 and SGK1 knock-down, but not AKT1/2 knock-down, recapitulate mitochondrial fission failure observed in *Rictor* depleted cells. (A)** Representative confocal images of NIH3T3 cells knocked-down for *Sgk1/2/3, Ndrg1* or *Akt1/2* and cultured in serum-free medium for 30 min in presence of MitoTracker green. Magnified insets are shown. Quantifications for mitochondrial number and mitochondrial morphology parameters are shown (36-71 cells from 3-5 independent experiments). **(B)** IB and quantifications for the indicated proteins in livers of 3-4 mo-old male mice injected with siRNAs against *Sgk1*, A*kt1/2* or *Ndrg1* and subjected to 14-16 h fasting (n=3-4). **(C)** AUC for OCR in si*Sgk1*, si*Akt1/2* or si*Ndrg1* livers of 14-16 h fasted mice (n=3-5). Ponceau is loading control. Individual replicates and means are shown. *P<0.05, **P<0.01, ***P<0.001 and ****P<0.0001. One-way ANOVA and Tukey's multiple comparisons test **(A** and **C)**; unpaired Student's t-test **(B)**. ns = not significant. Please refer to Table S2_statistical summary.

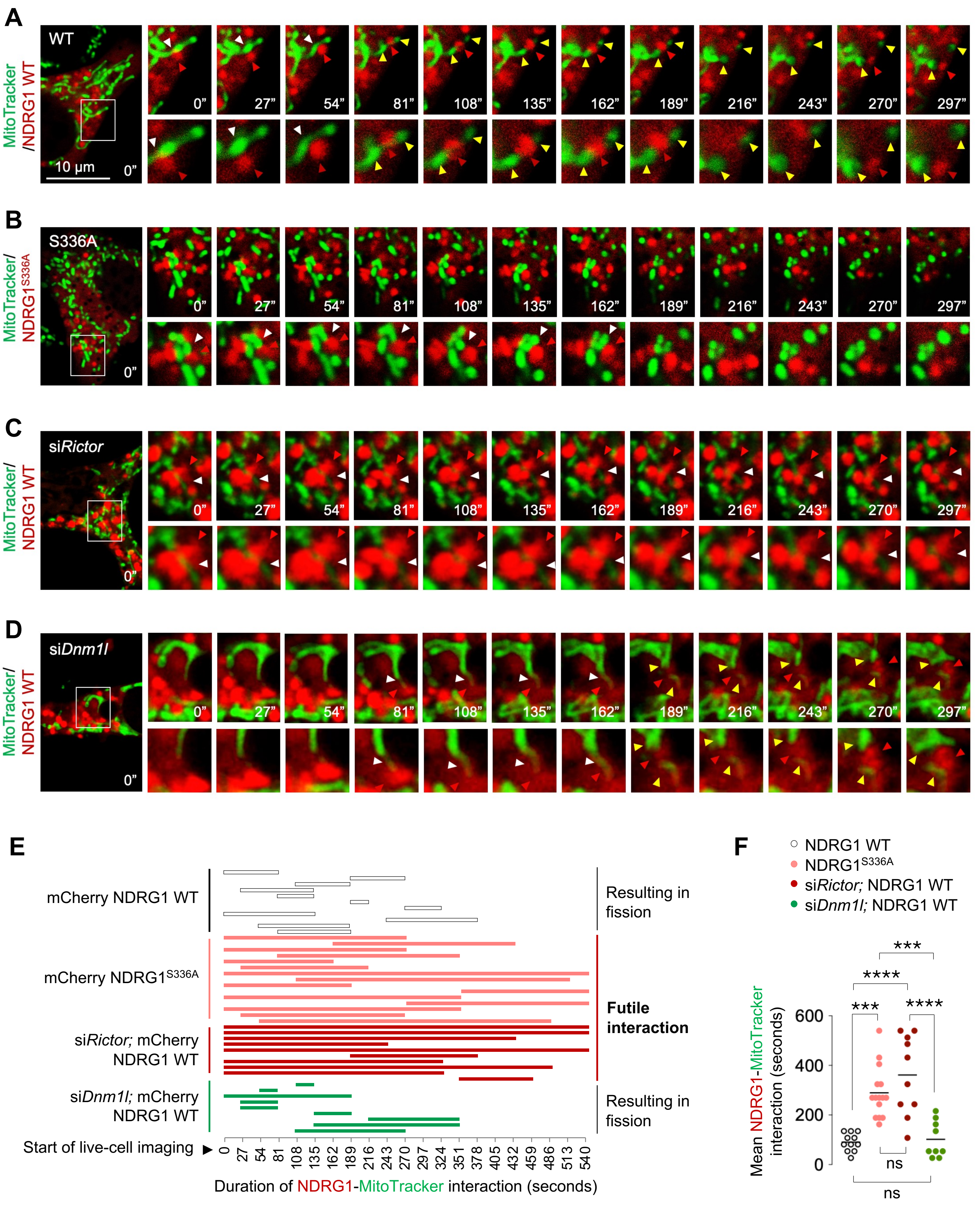
A
WT
MitoTracker/NDRG1 WT
10 μm
0"
27"
54"
81"
108"
135"
162"
189"
216"
243"
270"
297"
B
S336A
MitoTracker/NDRG1 S336A
C
siRictor
MitoTracker/NDRG1 WT
D
siDnm1l
MitoTracker/NDRG1 WT
E
mCherry NDRG1 WT
mCherry NDRG1 S336A
siRictor; mCherry NDRG1 WT
siDnm1l; mCherry NDRG1 WT
Start of live-cell imaging
Resulting in fission
Futile interaction
Resulting in fission
0 27 54 81 108 135 162 189 216 243 270 297 324 351 378 405 432 459 486 513 540
Duration of NDRG1-MitoTracker interaction (seconds)
F
NDRG1 WT
NDRG1 S336A
siRictor; NDRG1 WT
siDnm1l; NDRG1 WT
Mean NDRG1-MitoTracker interaction (seconds)
0
200
400
600
***
****
***
****
ns
ns

**Extended Data Fig. 11 | Scission of mitochondria by NDRG1 requires mTORC2 and NDRG1 phosphorylation at serine 336. (A-D)** Live cell imaging of mCherry-NDRG1 WT or mCherry-NDRG1$^{S336A}$ and MitoTracker green in siCon NIH3T3 cells or live cell imaging of mCherry-NDRG1 WT and MitoTracker in si*Rictor* or si*Dnm1l* NIH3T3 cells cultured in serum-free medium for 30 min. Red arrowheads: NDRG1. White arrowheads: NDRG1-mitochondria contact prior to fission. Yellow arrowheads: divided mitochondria after scission by NDRG1. Magnified insets are shown. **(E)** Quantification for duration/fate (fission vs. no fission) of interaction between mCherry-NDRG1 and mitochondria. A description of this quantitative approach is as follows: We quantified the duration of NDRG1-mitochondrial interaction events and whether each interaction led to mitochondrial fission (useful) or not (futile) as recorded via live cell imaging. The X-axis represents time in seconds—reflecting the duration of contact of NDRG1 with mitochondria; while individual-colored bars in the Y-axis represent the different cells/conditions. The length of each colored bar (when tracked from left to right) represents the time between the initiation of interaction of NDRG1 with mitochondria to the end of this interaction, which may or may not lead to mitochondrial division depending on the cell/condition. In addition to the time taken for mitochondrial fission by NDRG1, we noted if interaction with NDRG1 was **useful** (resulted in fission) or **futile** (no fission occurred). **(F)** Quantification for mean duration of mcherry-NDRG1/mitochondria (MitoTracker) interaction is shown. Individual replicates and means are shown. ***P<0.001 and ****P<0.0001. One-way ANOVA followed by Tukey's multiple comparisons test. ns = not significant. Please refer to Table S2_statistical summary.

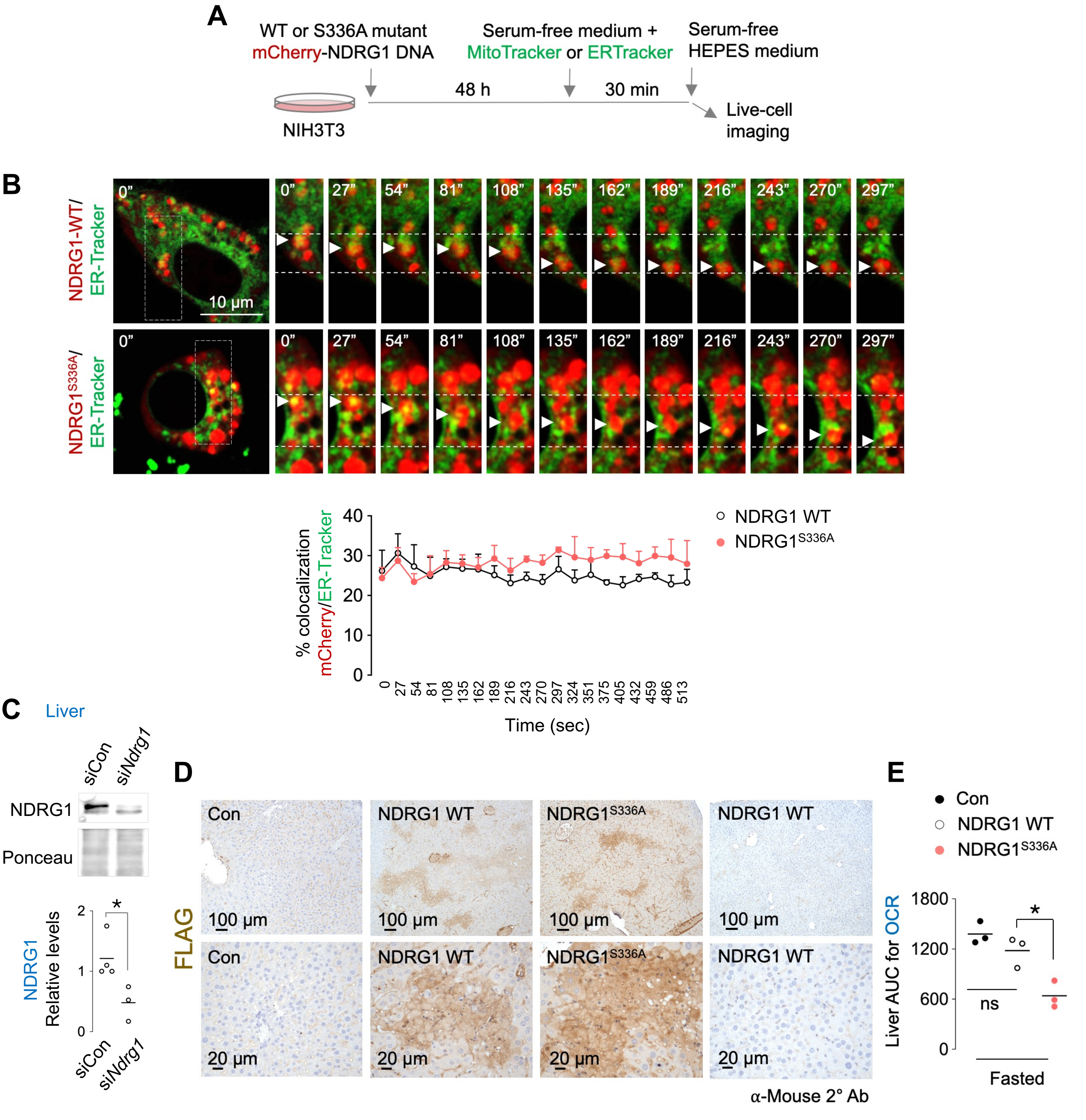


**Extended Data Fig. 12 | NDRG1 colocalizes with ER independent of its phosphorylation at Ser336, and demonstration of equivalent NDRG1 and NDRG1$^{S336A}$ in liver. (A)** Cartoon depicting experimental plan performed in **Fig. 3G** and **Extended Data Fig. 12B. (B)** Representative live-cell imaging of mCherry-NDRG1 WT or mCherry-NDRG1$^{S336A}$ and ER-Tracker green in NIH3T3 cells cultured in serum-free medium for 30 min. Magnified insets are shown. White arrowheads: NDRG1-ER contact. Quantification for % colocalization of mCherry with ER-tracker is shown. Values are mean ± SEM (n=3). **(C)** IB and quantification of NDRG1 in siCon or si*Ndrg1*-injected livers from mice subjected to 14-16 h fasting (n=3-4 mice). Ponceau is loading control. **(D)** Immunohistochemistry for equivalent FLAG expression in livers silenced for endogenous *Ndrg1* and then injected with Flag-tagged WT or S336A NDRG1 plasmid. 2° Ab-only control is shown. **(E)** Corresponding AUC for OCR is shown (n=3 mice). Individual replicates and means are shown. *P<0.05. 2-way ANOVA followed by Tukey's multiple comparison test **(B)**; Student's t-test **(C)**; One-way ANOVA followed by Tukey's multiple comparison test **(E).** Please refer to Table S2_statistical summary.

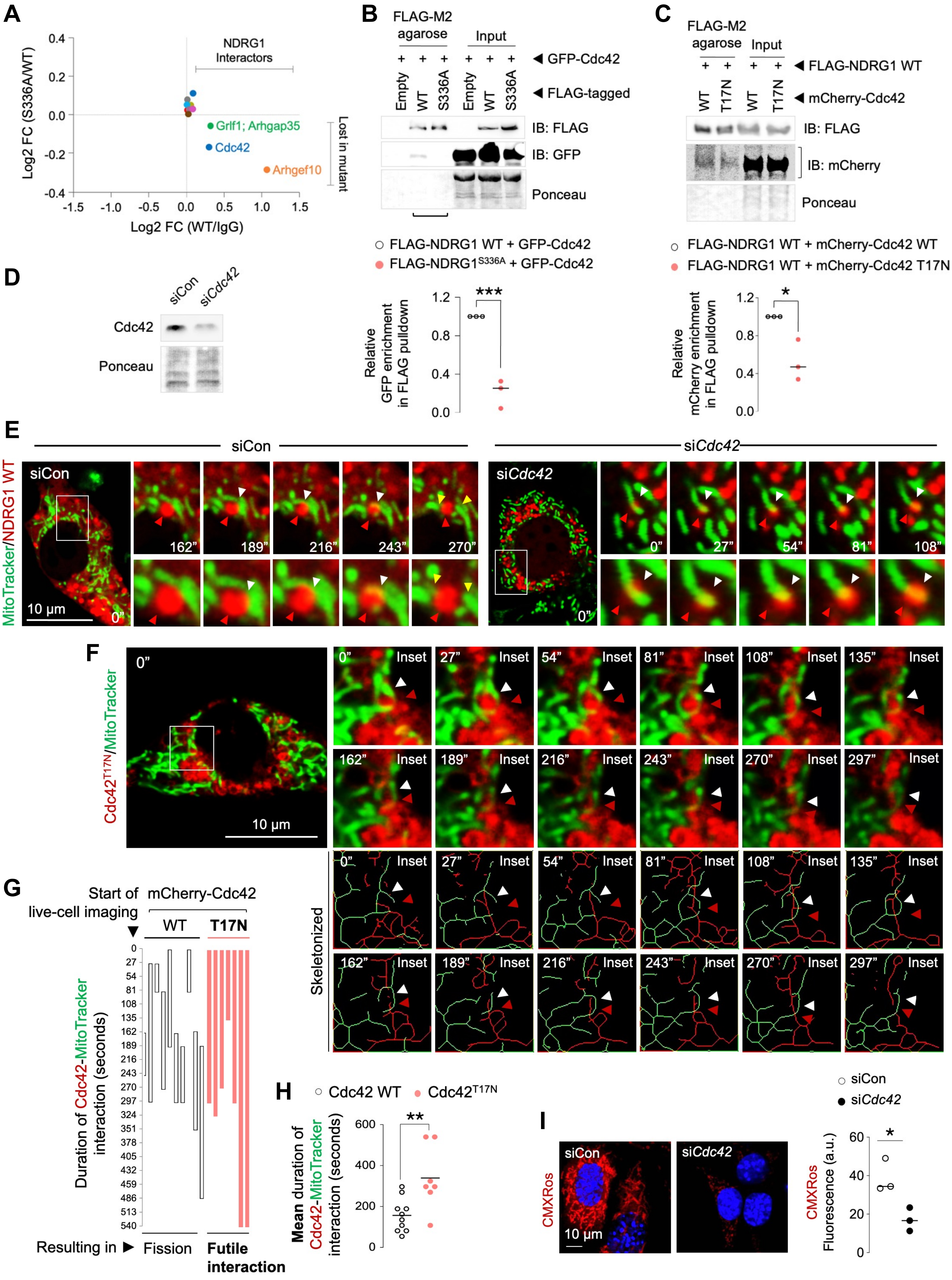

A
NDRG1 Interactors
Grlf1; Arhgap35
Cdc42
Arhgef10
Lost in mutant
Log2 FC (S336A/WT)
Log2 FC (WT/IgG)
B
FLAG-M2 agarose
Input
GFP-Cdc42
FLAG-tagged
Empty WT S336A Empty WT S336A
IB: FLAG
IB: GFP
Ponceau
FLAG-NDRG1 WT + GFP-Cdc42
FLAG-NDRG1S336A + GFP-Cdc42
Relative GFP enrichment in FLAG pulldown
***
C
FLAG-M2 agarose
Input
FLAG-NDRG1 WT
mCherry-Cdc42
WT T17N WT T17N
IB: FLAG
IB: mCherry
Ponceau
FLAG-NDRG1 WT + mCherry-Cdc42 WT
FLAG-NDRG1 WT + mCherry-Cdc42 T17N
Relative mCherry enrichment in FLAG pulldown
*
D
siCon siCdc42
Cdc42
Ponceau
E
siCon
siCdc42
MitoTracker/NDRG1 WT
10 µm
F
Cdc42T17N/MitoTracker
10 µm
Inset
Skeletonized
G
Start of live-cell imaging
mCherry-Cdc42
WT T17N
Duration of Cdc42-MitoTracker interaction (seconds)
Resulting in
Fission
Futile interaction
H
Cdc42 WT
Cdc42T17N
Mean duration of Cdc42-MitoTracker interaction (seconds)
**
I
siCon
siCdc42
CMXRos
10 µm
CMXRos Fluorescence (a.u.)
*

**Extended Data Fig. 13 | GTP bound-Cdc42 interacts with NDRG1 and supports mitochondrial fission. (A)** Log2 fold-change of interaction of NDRG1 WT or NDRG1$^{S336A}$ with Cdc42, Arhgap35 and Arhgef10 (n=3). **(B)** IB and quantification for GFP and FLAG levels in OA-treated (2.5 h) NIH3T3 cells co-expressing FLAG-NDRG1 WT or S336A with GFP-Cdc42 and subjected to pulldown of FLAG using FLAG M2 agarose (n=3). FLAG-tagged empty vector is the negative control. **(C)** IB and quantification for FLAG and mCherry in OA-treated (2.5 h) NIH3T3 cells co-expressing FLAG-NDRG1 WT and mCherry-Cdc42 WT or T17N mutant and subjected to pulldown of FLAG using FLAG M2 agarose (n=3). **(D)** IB for validation of *Cdc42* knockdown in NIH3T3 cells. **(E)** Live-cell imaging of mCherry-NDRG1 WT and MitoTracker green in siCon or si*Cdc42* NIH3T3 cells. Magnified insets are shown. Red arrowhead: NDRG1 mediating fission. White arrowhead: NDRG1-mitochondrial colocalization before fission. Yellow arrowheads: divided mitochondria after fission. **(F)** Representative live-cell imaging in cells expressing dominant negative mcherry-Cdc42$^{T17N}$ mutant in presence of MitoTracker green to label mitochondria. Red arrowheads: Cdc42. White arrowheads: Cdc42-mitochondria contact prior to fission. Magnified insets are shown. **(G)** Graphical representation for duration of interaction between mCherry-Cdc42 WT or Cdc42$^{T17N}$ and mitochondria (Mitotracker), and whether interactions lead to division or are futile. **(H)** Quantification for mean duration of interaction (n=7-10 cells from 3-4 independent experiments). **(I)** MitoTracker CMXRos fluorescence in siCon and si*Cdc42* cells cultured in serum-free medium in the presence of OA for 5 h (102-116 cells from 3 independent experiments). Ponceau is loading control. Individual replicates and means are shown. *P<0.05, **P<0.01 and ***P<0.001. Unpaired Student's t-test. Please refer to Table S2_statistical summary. Please refer to excel files, Proteomics5_FLAG_NDRG1_coIP.

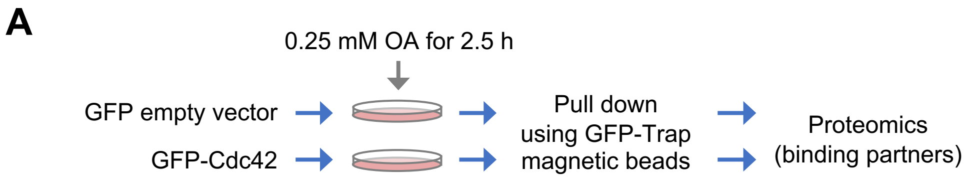


**B**

| Position | Accession | Description | Gene Symbol | Fold Change (GFP-CDC42/GFP) | P-value | Score |
|---|---|---|---|---|---|---|
| 4 | Q9JM96 | Cdc42 effector protein 4 | *Cdc42ep4* | 5.96 | 8.51 | 50.71 |
| 5 | Q91W92 | Cdc42 effector protein 1 | *Cdc42ep1* | 6.08 | 7.31 | 44.46 |
| 6 | Q9JKF1 | Ras GTPase-activating-like protein IQGAP1 | *Iqgap1* | 7.92 | 6.61 | 52.36 |
| 7 | Q99PT1 | Rho GDP-dissociation inhibitor 1 | *Arhgdia* | 6.39 | 6.33 | 40.42 |
| 12 | Q8JZX9 | Cdc42 effector protein 2 | *Cdc42ep2* | 4.36 | 5.40 | 23.54 |
| 22 | Q9JI08 | Bridging integrator 3 | *Bin3* | 3.77 | 3.94 | 14.83 |
| 24 | Q9DAK3 | Rho-related BTB domain-containing protein 1 | *Rhobtb1* | 6.61 | 3.90 | 25.78 |
| 40 | Q61599 | Rho GDP-dissociation inhibitor 2 | *Arhgdib* | 4.98 | 3.27 | 16.28 |

Significantly enriched in Cdc4 pull downs (positions 4–12)

***Cut off (P-value 4.32)***

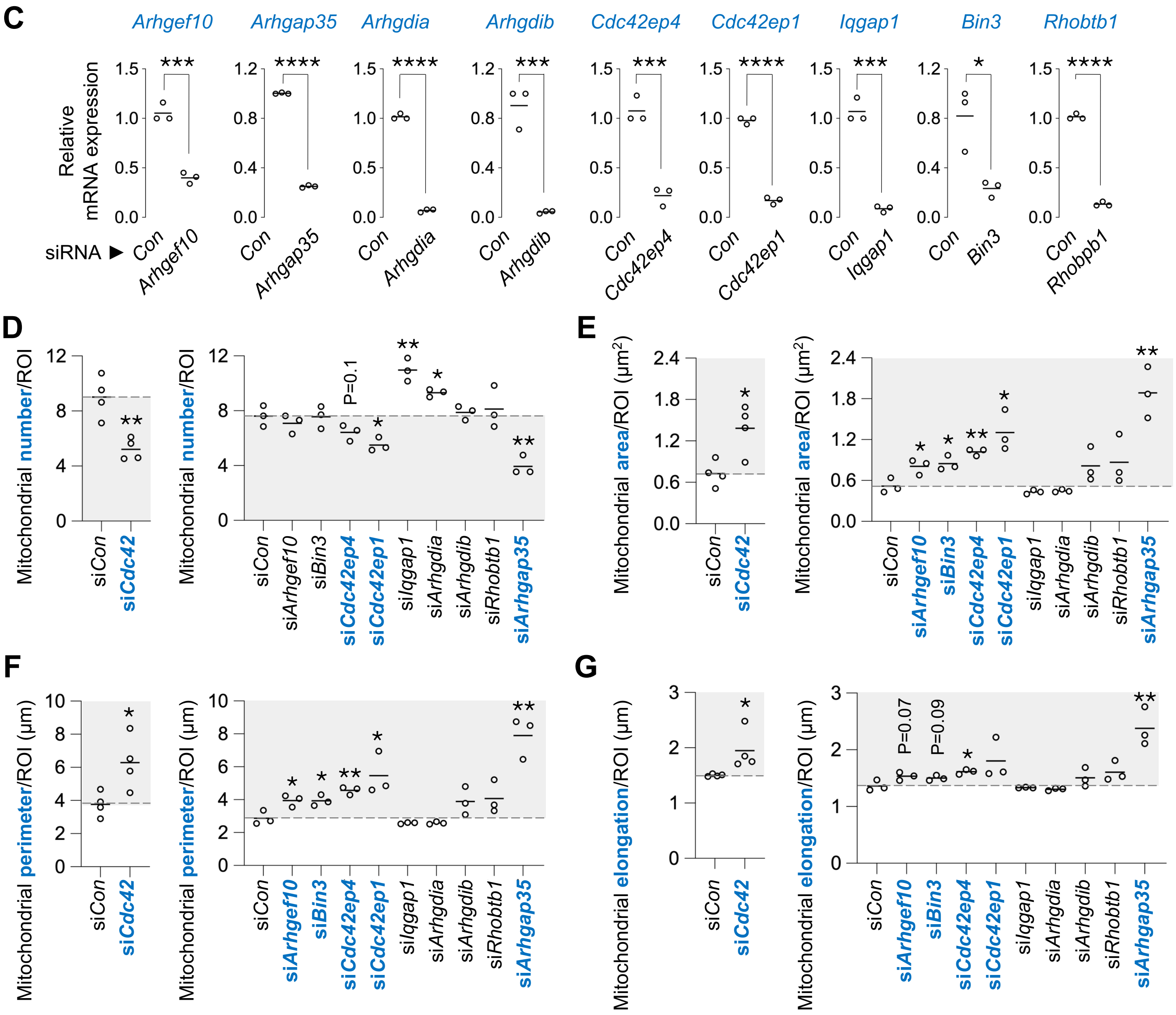

**Extended Data Fig. 14 | An siRNA screen to target interacting partners of Cdc42 reveals effectors and regulators controlling mitochondrial dynamics.** **(A)** Cartoon depicting the experimental plan to pulldown GFP-Cdc42 and identify interacting partners via proteomics. Empty GFP vector served as control. **(B)** Significantly enriched interacting partners of Cdc42 that belong to the Rho family of GTPases. A P-value cut-off of 4.32 was used as threshold. **(C)** qPCR in NIH3T3 cells to validate the silencing of the selected Cdc42-binding partners identified by proteomics (n=3). Quantifications for **(D)** mitochondrial number, **(E)** mitochondrial area, **(F)** mitochondrial perimeter and **(G)** mitochondrial elongation (n=37-40 cells from 3 independent experiments). Gray areas indicate mitochondria fission-deficient models. Individual replicates and means are shown. *$P<0.05$, **$P<0.01$, ***$P<0.001$ and ****$P<0.0001$, Student's t-test. Please refer to Table S2_statistical summary. Please refer to excel files, Proteomics6_GFP_Cdc42_coIP.

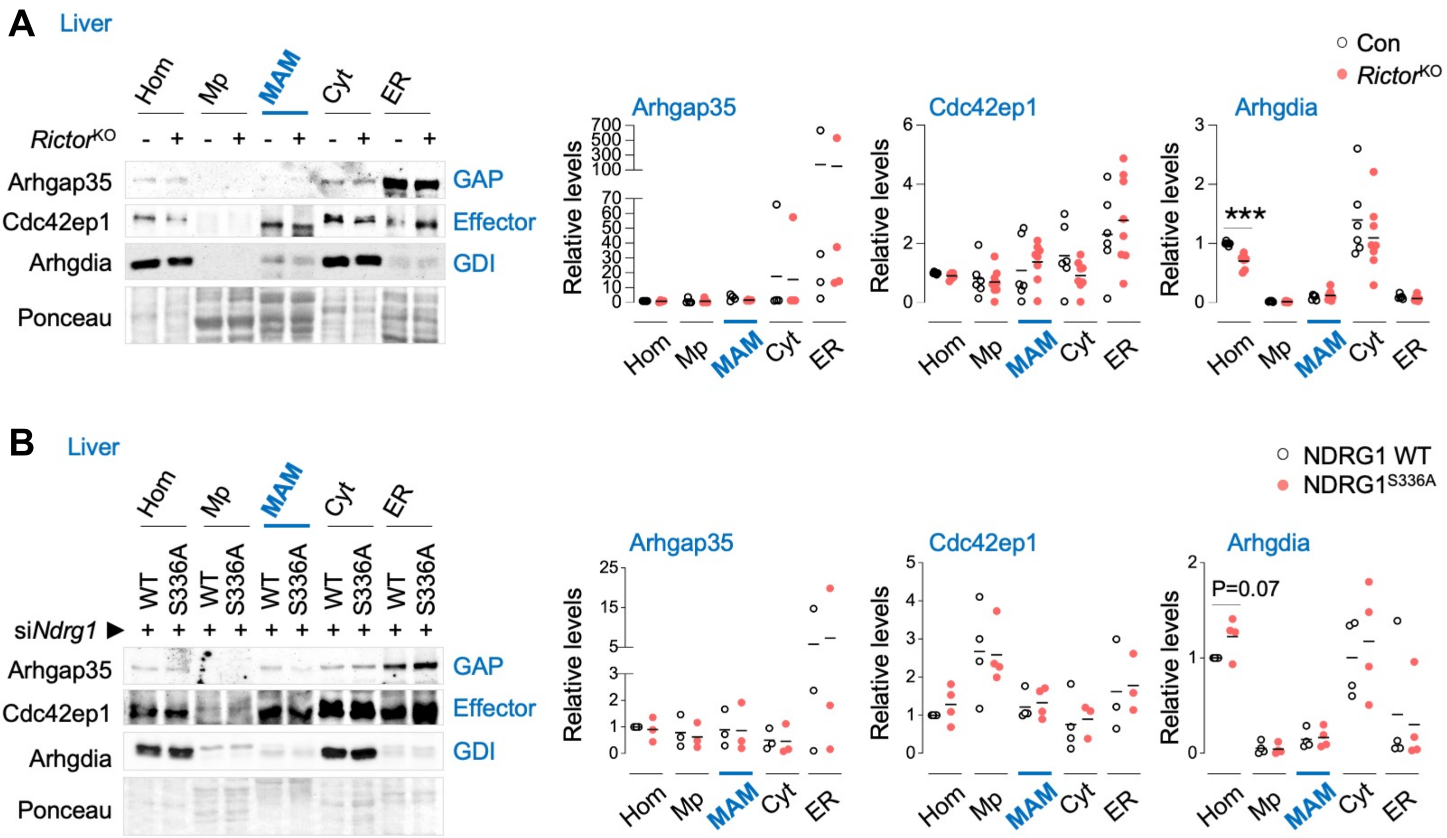


**Extended Data Fig. 15 | Cdc42 regulatory proteins are enriched in MAMs. (A, B)** IB and quantification for indicated proteins in Hom, Mp, MAMs, Cyt, and ER fractions from livers of **(A)** 5-6 mo-old Con or *Rictor*$^{KO}$ male mice (n=3-5 mice), and **(B)** 3-4 mo-old NDRG1 WT or NDRG1$^{S336A}$ male mice co-injected with siRNA against endogenous *Ndrg1* and fasted for 14-16 h (n=3-4 mice). Ponceau is loading control. Individual replicates and means are shown. ***P<0.001, unpaired Student's t-test. GAP: GTPase activating protein; GDI: GDP dissociation inhibitors. Please refer to Table S2_statistical summary.
